# Development of a recombineering system for the acetogen *Eubacterium limosum* with Cas9 counterselection for markerless genome engineering

**DOI:** 10.1101/2024.04.09.588731

**Authors:** Patrick A. Sanford, Benjamin M. Woolston

## Abstract

*Eubacterium limosum* is a Clostridial acetogen that efficiently utilizes a wide range of single-carbon substrates and contributes to metabolism of health-associated compounds in the human gut microbiota. These traits have led to interest in developing it as a platform for sustainable CO_2_-based biofuel production to combat carbon emissions, and for exploring the importance of the microbiota in human health. However, synthetic biology and metabolic engineering in *E. limosum* have been hindered by the inability to rapidly make precise genomic modifications. Here, we screened a diverse library of recombinase proteins to develop a highly efficient oligonucleotide-based recombineering system based on the viral recombinase RecT. Following optimization, the system is capable of catalyzing ssDNA recombination at an efficiency of up to 2%. Addition of a Cas9 counterselection system allows recombination to reach an efficiency of up to 100%, enabling creation of genomic point mutations in a scarless and markerless manner. We deployed this system to create a clean knockout of the extracellular polymeric substance (EPS) gene cluster, generating a strain incapable of biofilm formation. This approach is rapid and simple, not requiring laborious homology arm cloning, and can readily be retargeted to almost any genomic locus. This work overcomes a major bottleneck *Eubacterium limosum* genetic engineering by enabling precise genomic modifications, and provides both a roadmap and associated recombinase plasmid library for developing similar systems in other Clostridia of interest.

**Graphical Abstract:** 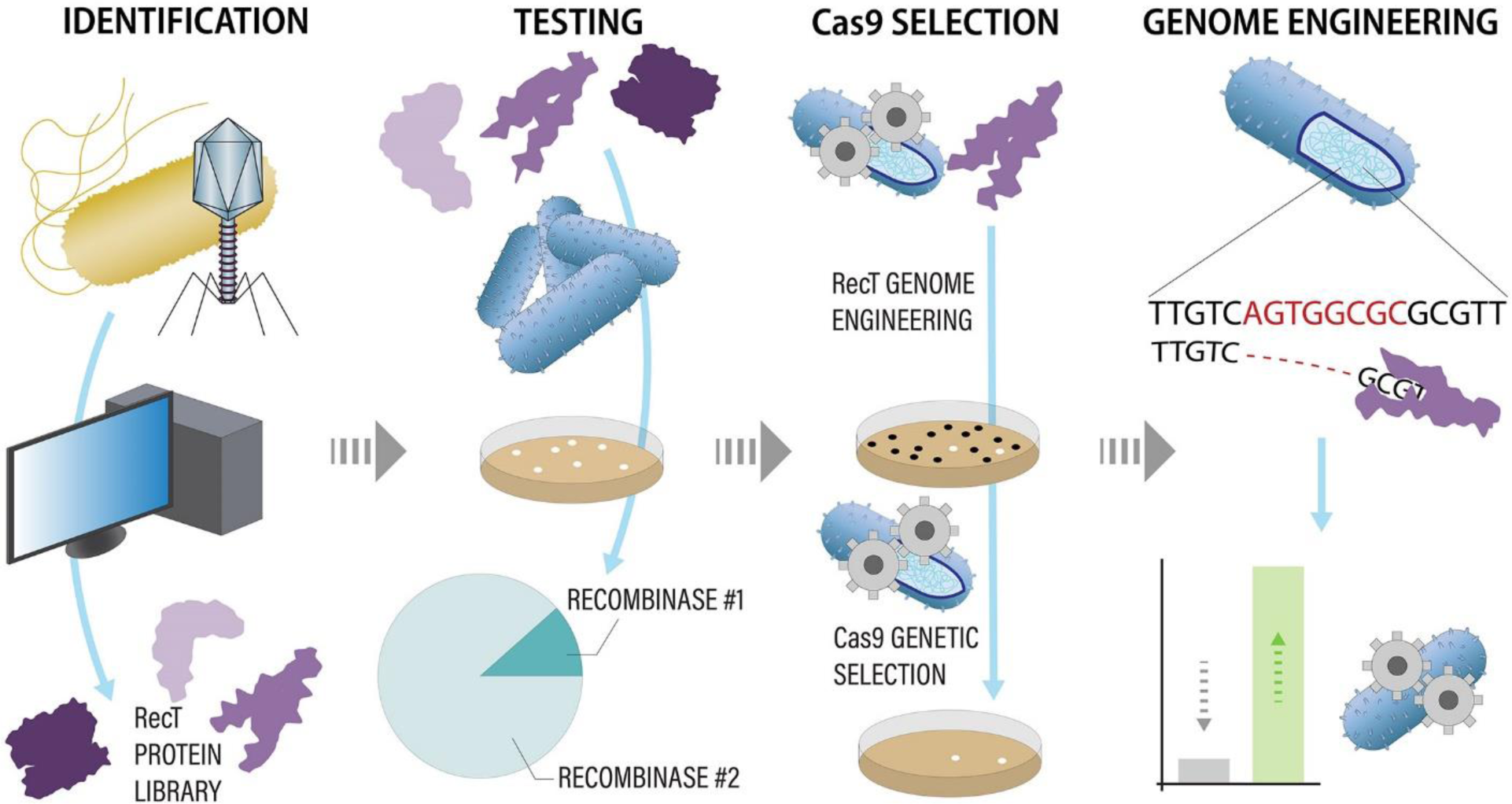

## Introduction

*Eubacterium limosum* is an acetogenic bacterium of growing interest thanks to its metabolic flexibility and role as a member of the human gut microbiota [1], [2]. Acetogens, including *E. limosum*, are capable of metabolizing single carbon substrates such as carbon dioxide at very high efficiency, upcycling them into value added products, providing an attractive avenue for greenhouse gas sequestration in processes which are currently being commercialized [3]–[6]. *E. limosum* is particularly suited to such applications due to its ability to metabolize an expanded range of substrates including methanol and formate [7]. Additionally, as a member of the gut microbiota, it has been found to demethylate a range of dietary compounds of relevance to human health, including flavonoids and carnitine, making it an organism of interest for studying microbiota-related pathologies [8], [9]. Amongst acetogens, *E. limosum* possesses a growing engineering toolbox including an analyzed genome and transcriptome, an expanded range of selection markers, and libraries of genetic elements, all of which have been used primarily in plasmid-based engineering applications [5], [10]–[13]. Recently, more additions have been made to the organism’s genome engineering repertoire including a CRISPR/dCas9 system for gene silencing and a toxin/antitoxin system for markerless knockout generation [14], [15]. However, the organism’s engineering toolbox could benefit from rapid genome engineering techniques such as homologous recombination to further leverage its native strengths. We recently reported utilizing the microbe’s native RecA-based recombination machinery to perform genomic knock-ins and targeted gene deletions [16]. However, this system requires tedious cloning, is only capable of catalyzing recombination with very long overhangs, at very low efficiency, and with the integration of a selectable marker, making it unsuitable for high-throughput engineering, creation of point mutations, or markerless knockouts. This is especially relevant since, as an acetogen, *E. limosum* lacks the non-homologous end joining DNA repair pathway, further limiting its genome engineering capacity [17]. The development of more recombineering systems has therefore been of growing interest in the acetogen synthetic biology community, as development of such tools would significantly streamline engineering efforts in these organisms [18], [19]. Without these tools, engineering is limited to large edits, complex multi-step workflows, and the integration of selection markers, rendering interrogation of specific genomic point mutations or construction of complex genetic circuits impossible.

Homologous recombination (HR) in bacteria via phage-derived recombinases is one of the fundamental tools in the synthetic biology toolbox [20]. The most well-known examples of this are the lambda Red (in which the recombinase enzyme is Beta) and RecT (in which the recombinase enzyme is RecT) recombineering systems in *Escherichia coli*, which enable gene modification via point or large mutations, gene knockout, and random oligonucleotide (oligo) mediated mutagenesis. These tools have significantly contributed to *E. coli*’s development as the de facto bacterial engineering host [21]–[23]. More recently, Cas9 has been added to these recombineering systems as a ‘proof-reading’ counterselection agent, cleaving unmutated DNA and thereby boosting recombineering efficiency to 100%, often in a scarless and markerless manner [24], [25]. While Beta from lambda Red is the most commonly used recombinase in *E. coli*, this protein family is not widespread and has limited host range [26], [27]. Across all bacteria, recombinases from the phage protein superfamily RecT are the most broadly distributed [28]. RecT binds to single strand DNA and anneals it to homologous ssDNA in the host during DNA replication in a process independent of native RecA-mediated DNA repair [29], [30]. Given their functional orthogonality to native DNA repair, it has been hypothesized that RecT recombinase may possess much broader host range than Beta, unconstrained by taxonomy. Indeed, a recent screen of a large library of RecT proteins in *E. coli* revealed a surprising lack of correlation between recombinase activity and phylogenetic similarity [27]. Consequently, RecT recombinases are a promising family to investigate for implementing recombineering in underdeveloped microbes. Here, to enhance the synthetic biology capacity of *E. limosum*, we developed a highly efficient RecT-based recombineering system with Cas9 counterselection for markerless genome engineering.

## Results and Discussion

### Recombinase Library Screening

Given the unclear importance of the phylogenetic similarity between recombinase source and target host, we assembled a library of 57 unique recombinases for assessment in *E. limosum* selected from a broad range of sources including both characterized and putative enzymes [27], [28] (**Table S1**). We began constructing the recombinase library by selecting a group of published and/or characterized recombinases including GP35 from *Bacillus subtilis*, Beta from *E. coli*, and RecTs from *Clostridium perfringens*, *Pseudomonas syringae*, and *E. coli* [27], [28], [31], [32]. Next, since strong bacterial recombinases typically originate from phages, we utilized the PHASTER tool to search *E. limosum* strains ATCC 8486, SA11, and 81C1 for prophage regions [33]. This analysis revealed several prophages within SA11 and 81C1 containing genes annotated as putative *recT*s, which we selected for testing. Following on from this analysis, PHASTER also identified the original phages likely responsible for these infections, from which we identified the original *recT* genes and added them to the library. These were checked for genes annotated as *recT* as inferred from homology, of which, across the three *E. limosum* strains, phages PLE3, phiCT19406A, phiCT19406B, phiCTC2A, phiCTC2B, and phiCT19406C encoded suitable genes. These additional phage enzymes were added because prophages within bacterial genomes are commonly inactivated via a plethora of mechanisms including point mutations, regulatory deactivation, and gene shuffling including operon inversion, among others, making it possible for a prophage-derived RecT to be non-functional [34].

Next, we searched for recombinase genes in related organisms in the NCBI genome database, screening the class Clostridia and phylum Firmicutes in alignment with *E. limosum’s* taxonomy. From this screen, we selected seven additional proteins annotated as RecT. Then, based on recent data indicating that phylogenetic similarity may not be a factor in recombinase performance, we expanded our criteria and selected 22 additional *recT* genes from a broad range of bacterial backgrounds [27]. Finally, we applied the same approach to phages and selected 12 *recT* genes from a diverse range of bacteriophages. The 57 recombinase genes were refactored for expression in *E. limosum* and synthesized into plasmid-borne expression cassettes under the control of the inducible p2TetO1 promoter [13], [35].

We assessed recombinase performance in a fashion similar to procedures reported previously by creating a base ‘recombinase test strain’ (RTS) encoding a mutated genomic copy of *ermB* (*ermB’*) for inactivated clarithromycin resistance with three stop codons inserted early in the sequence to prevent expression [27], [31] (**Figure S1**). We transformed this strain individually with each of the recombinase expression constructs, then pooled the resulting strains into a single library culture (**Table S2**). We transformed the library culture with 120 bp recombinogenic oligos targeting the stop codons in *ermB’* to return them to the original coding sequence and restore antibiotic resistance function (**Figure 1a, Table S3**). This initial library test was performed with oligos in both the sense and antisense orientations as well as on the RTS strain without a RecT recombinase to assess the efficacy of the native RecA recombinase in catalyzing oligo-length ssDNA recombination. While it is generally accepted that recombinases function at much higher efficiency annealing heterologous ssDNA to the lagging strand of genomic DNA during DNA replication, it is possible for them to anneal foreign ssDNA to the leading strand as well [20]. So, while antisense ssDNA oligos are typically used for recombineering, we included sense orientation oligos as well as this orientation has occasionally been known to recombine [36].

**Figure 1:**
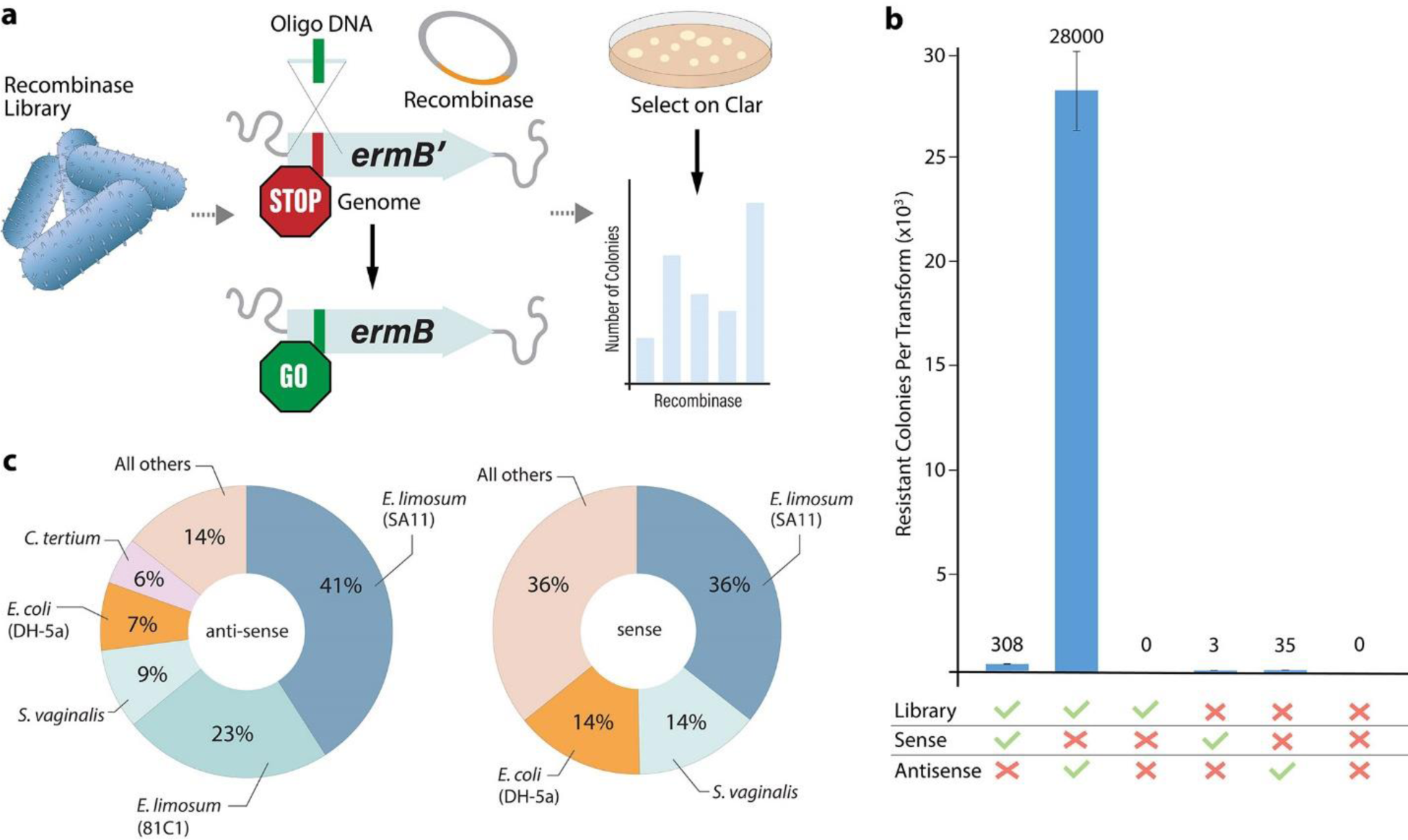
Screening a recombinase library reveals a highly active RecT protein from the *E. limosum* SA11 prophage. a) Scheme for testing recombinase library by activating a mutated *ermB* gene containing early stop codons. b) Number of clarithromycin-resistant colonies obtained by transforming recombinase library or no-library control with 120 bp ssDNA recombinogenic oligos. Error bars indicate standard deviation of biological triplicate transformations. c) Proportion of clarithromycin resistant colonies encoding any given recombinase from conditions treated with either sense or antisense oligos for *ermB’* correction.

These transformations revealed that the library possessed significant recombinase function with a strong preference for antisense ssDNA, resulting in about 28,000 recombined colonies for antisense and 308 for sense per transformation (**Figure 1b**). Additionally, the native RecA recombinase is capable of catalyzing some very low level of recombination with ssDNA oligos, but at an efficiency many orders of magnitude below the RecT recombinases. Sequencing of resistant colonies showed that plasmid pEL11, encoding the RecT from the *E. limosum* SA11 prophage, was the most abundant library member, being responsible for 41% of recombined colonies with antisense ssDNA and 36% with sense ssDNA (**Figure 1c**). Based on these findings, we selected SA11 RecT and antisense ssDNA as our recombinase and DNA orientation for further optimization.

The difference in efficiency between sense and antisense DNA is perhaps unsurprising given the high availability of the lagging strand in the genomic DNA during replication and that this phenomenon is observed across other recombinases [20], [37]. While orientation specificity was as expected, the distribution of recombinases is interesting in that it seems to indicate that phylogenetic similarity between source and host is an important factor for recombinase function in *E. limosum*, which is in contrast to recent results in *E. coli* [27]. In fact, the two most efficient recombinases in *E. limosum* ATCC 8486 were sourced from prophages in the genomes of two other strains of *E. limosum*: SA11 and 81C1, accounting for nearly two-thirds of all recombined colonies. RecT from another closely related source, *Clostridium tertium*, also contributed significantly, and ultimately about 73% of all recombined colonies possessed *recT* genes sourced from the class Clostridia. However, this is not to say that source similarity is the only determinant of recombinase function for *E. limosum*, as recombinases isolated from distantly related *Sneathia vaginalis* and *E. coli* both catalyzed efficient recombination. Surprisingly, the prophage from SA11 which encoded the best RecT enzyme originated from lactobacillus phage PLE3, a phage whose typical host is not closely related to *E. limosum*. This finding was unexpected and adds more complexity to the question of whether phylogenetic similarity is important in recombinase selection given this phage’s normal host range.

### Recombineering System Optimization

Recombination efficiency has previously been optimized in other organisms by a variety of means including varying DNA concentration and overhang length, modifying DNA structure, mutating the recombinase enzyme, supplementing the reaction with small-molecule enhancers, and changing recovery and growth conditions [38]–[40]. Total homology length is an essential consideration in recombinase function, as some recombinases work best with specific lengths of homology between heterologous DNA and genomic template [38]. Additionally, the concentration at which the heterologous DNA is provided is an important consideration as this is known to have a significant impact on transformation efficiency in general [41]. Therefore, we began optimizing our system by assessing the impact of varying oligo lengths and concentrations utilizing 120, 60, and 30 bp oligos spanning a final concentration of 1-1000 ng/μL in the transformation reaction. Maximum recombination efficiency was achieved using 60 bp oligos at a concentration of 1000 ng/μL (**Figure 2a**). We were unable to examine any higher DNA concentrations due to arcing during electroporation. Interestingly, we obtained very few colonies for any concentration of 30 bp oligo DNA. These oligos encode a total of 26 bp homology with the host gDNA, which implies that this length is insufficient for recombination catalyzed by RecT. However, maximum efficiency was observed with the 60 bp oligos encoding a total homology of 56 bp, which indicates that the minimum homology length for recombination is between 26 and 56 bp. Additionally, increasing the oligo length beyond 60 bp did not significantly improve efficiency for any concentration, however, efficiency did significantly decrease with the 120 bp oligos at the highest concentration of 1000 ng/μL.

**Figure 2:**
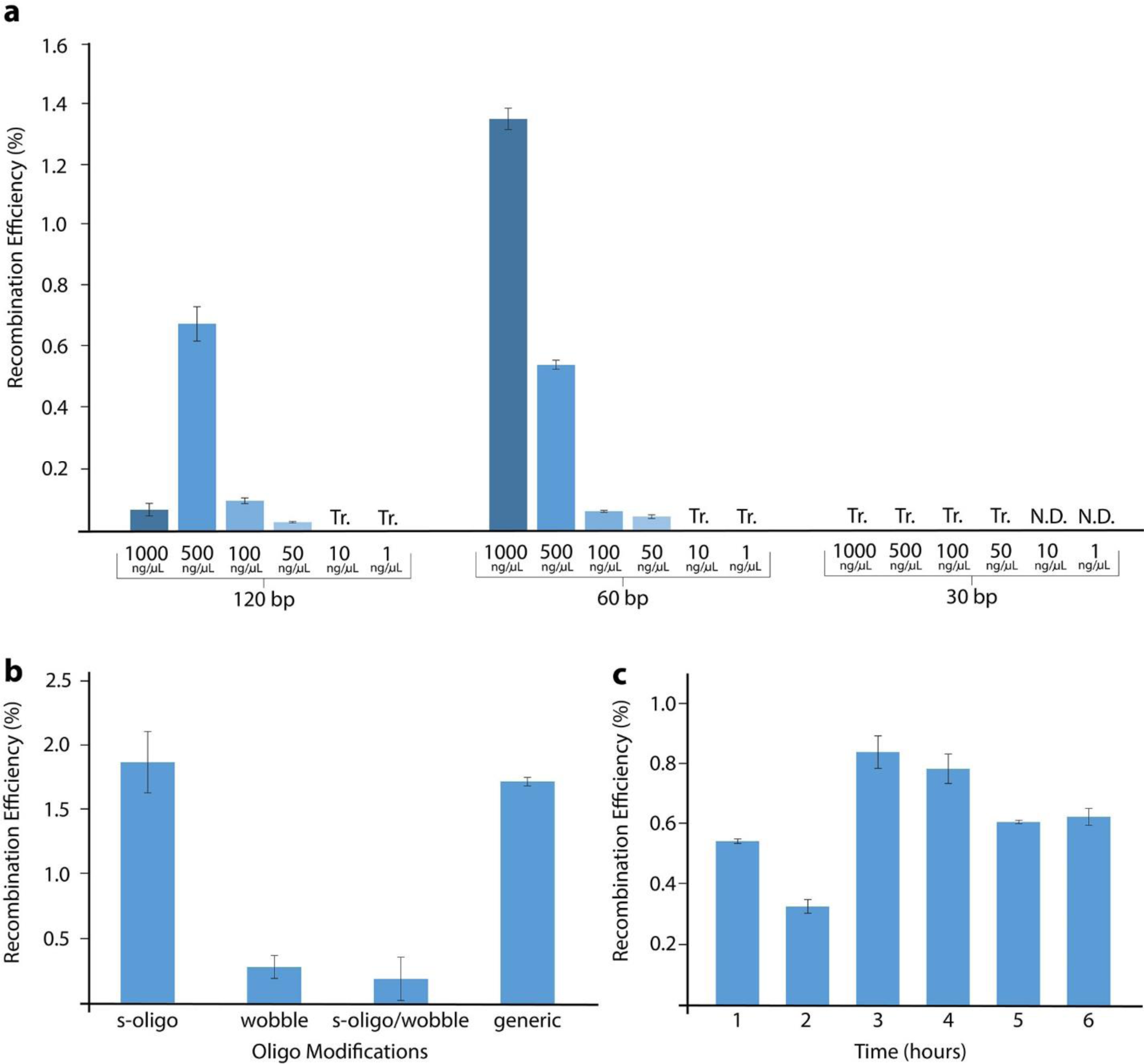
Oligo and process modifications affect recombineering efficiency. a) Effect of oligo size and concentration on recombineering efficiency (% viable cells recombined). B) Effect of oligo modifications on recombineering efficiency. ‘S-oligo’ refers to oligos with phosphonothioate bases in the first and final three positions of the oligo, ‘wobble’ refers to oligos encoding consecutive wobble-base codon mutations in addition to the original targeted mutation, and ‘generic’ refers to oligos without either of these alterations. c) Effect of recovery time on recombineering efficiency. In all cases error bars represent standard deviation of biological triplicate transformations. N.D: not detected, Tr: trace.

Next, we used the optimized conditions of 60 bp oligos at a concentration of 1000 ng/μL to assess the impact of oligo modifications on recombination efficiency by two strategies: mutating consecutive wobble-bases, and preventing exonuclease function from reducing the effective concentration of recombinogenic DNA [38], [42]. It has previously been reported for recombineering systems in other organisms that efficiency can be improved by mutating consecutive wobble-bases, a modification which is hypothesized to enable evasion of native DNA proofreading machinery [38]. It has further been shown that oligo degradation by exonucleases can reduce recombination efficiency by diminishing the effective concentration of recombinogenic DNA [42]. We obtained a panel of oligos encoding mutations to consecutive wobble bases in addition to the repair for *ermB’*, and also obtained oligos with phosphorothioate bases in the first and final three positions to minimize exonuclease degradation [43] (**Table S3**). Contrary to expectations, these modifications had no beneficial effect on recombination efficiency, and in fact the additional wobble base mutations reduced efficiency (**Figure 2b**). This may indicate that, in *E.* limosum, the potential enhancing effect of evading mismatch repair is outcompeted by a reduced efficiency of recombination. It is also likely that the concentration of 1000 ng/μL is so high that it overwhelms the cell’s exonucleases, rendering the phosphorothioate modification unnecessary.

Finally, we assessed the effect of recovery time on recombination efficiency, since it has been hypothesized that differences in cell survival and growth between recombined and unrecombined strains following transformation affect overall transformation efficiency observed after plating for single colonies [38]. We tested a panel of recovery times spanning 1-6 hours and observed only minor differences across that range, with efficiency relatively stable from 3 hours onwards (**Figure 2c**). Optimal efficiency was observed in a 3-6 hour recovery time window, which is consistent with the optimal recovery time for general transformations in *E. limosum* [13]. Using optimized conditions, the recombineering system is capable of ∼2% recombineering efficiency (1/50 cells possess the desired mutation from recombinase activity), which in most cases equates to more than 2×10^6^ recombined colonies per transformation. This should be sufficient for library-based engineering approaches given careful screening criteria, however, in most cases, it is not sufficient for markerless genome engineering.

### Implementation of Cas9 Counterselection

To enable high efficiency markerless genome engineering, we combined our recombineering system with Cas9 counterselection, a strategy which has enabled near perfect recombineering efficiency in *E. coli* [44]. We tied the recombinase-mediated mutation to the deletion of a PAM site and in tandem expressed Cas9 with an sgRNA targeting the mutation (**Table S4**). Since the PAM site is essential for Cas9 activity, the Cas9 enzyme will only bind and cleave unrecombined genomes and therefore only correctly recombined strains will survive Cas9 expression. Additionally, a second sgRNA was included to target the recombinase plasmid to reduce the probability of aberrant recombination after obtaining the desired mutation (**Figure 3**). To demonstrate the applicability of this method in *E. limosum*, we used a strain with the colorimetric reporter gene *bgaB* integrated into the genome (**Figure S2**). In the presence of S-Gal and ammonium ferric citrate, cells expressing the β-galactosidase enzyme BgaB produce an insoluble black product which makes them easily differentiated by visual screening [13]. Analysis of a homology model of BgaB identified Glu-303 as the catalytic nucleophile (**Figure S3**), consistent with Glu-537 in *E. coli* β-galactosidase which, when mutated, abolishes activity of that enzyme [45]. To eliminate function of BgaB we designed a recombinogenic oligo, oPAS518, to mutate this codon to instead code for alanine (**Table S3**) [46]. The oligo additionally possessed a noncoding mutation to base 912 of the *bgaB* gene from guanine to adenine to remove the PAM site adjacent to the coding mutation.

**Figure 3:**
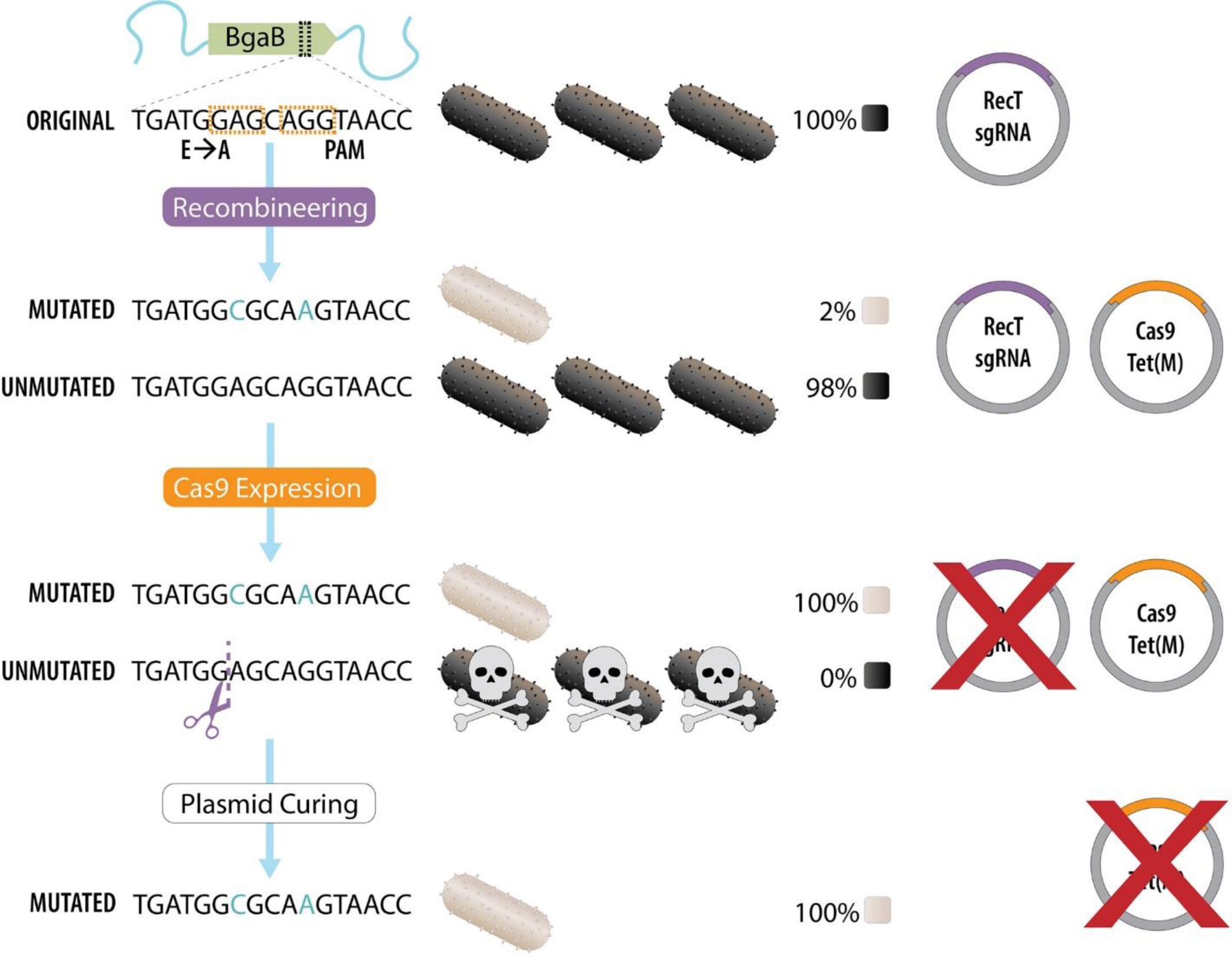
Scheme for improved recombineering efficiency using Cas9 counterselection. Plasmid encoding RecT and two sgRNAs – one targeting a PAM site near the desired mutation and one targeting the RecT plasmid – is transformed into *E. limosum* before that strain is again transformed with Cas9 plasmid and recombinogenic oligos and selected on tetracycline. Expression of Cas9 kills unrecombined colonies and destroys the recombinase plasmid. Finally, surviving colonies are sub-cultured to cure the Cas9 plasmid resulting in a strain with a markerless and scarless genomic point mutation.

We constructed plasmid pEL11-BgaB encoding RecT, inducibly expressed from the p2TetO1 promoter, along with two sgRNAs: one targeting the plasmid itself, and the other targeting *bgaB* in the genome **(Tables S3 – S4)** [13], [35]. We transformed this plasmid into an *E. limosum* strain with genomic *bgaB*, and prepared competent cells with RecT induced during growth. We transformed this strain with the Cas9-bearing plasmid along with the recombinogenic oligo encoding the E303A BgaB mutation and selected on tetracycline plates. Since the Cas9 plasmid, pRec-Cas2, encodes tetracycline resistance, only cells which received the Cas9 plasmid survived plating. Consequently, these cells also received the Cas9 counterselection, killing those which were not recombined by RecT, leaving only properly recombined cells. Following transformation, we picked colonies from each condition and treated them with S-Gal and ammonium ferric citrate to assess BgaB activity. The experimental condition with RecT, the recombinogenic oligo, and Cas9 counterselection resulted in all colonies containing inactive BgaB as evidenced by their uniform white color. In contrast, control transformations lacking either RecT, oligos, or Cas9, retained their black phenotype (**Figure 4a, b**). These results were also confirmed by DNA sequencing which showed white colonies possessed the two targeted point mutations, while black colonies retained the unmutated sequence (**Figure S4**). Additionally, the absence of colonies on tetracycline plates for conditions lacking the recombinogenic oligo but transformed with Cas9 plasmid encoding *Tet(M)* confirms that Cas9 counterselection killed unrecombined colonies, with no escapees (**Figure 4c**). While it is sometimes possible for escapees to evade Cas9 counterselection, we did not observe the phenomenon here. It is interesting to note that the number of colonies observed on tetracycline plates for the experimental condition treated with RecT, Cas9, and recombinogenic oligos was far lower than would normally be expected for plasmid transformations. In this strategy, a single cell must receive both the plasmid and recombinogenic oligo during the transformation in order to obtain the mutation and survive on the selection plate. Receiving either one of these is already unlikely, and the necessity of receiving both compounds this unlikeliness, resulting in far fewer correct colonies.

**Figure 4:**
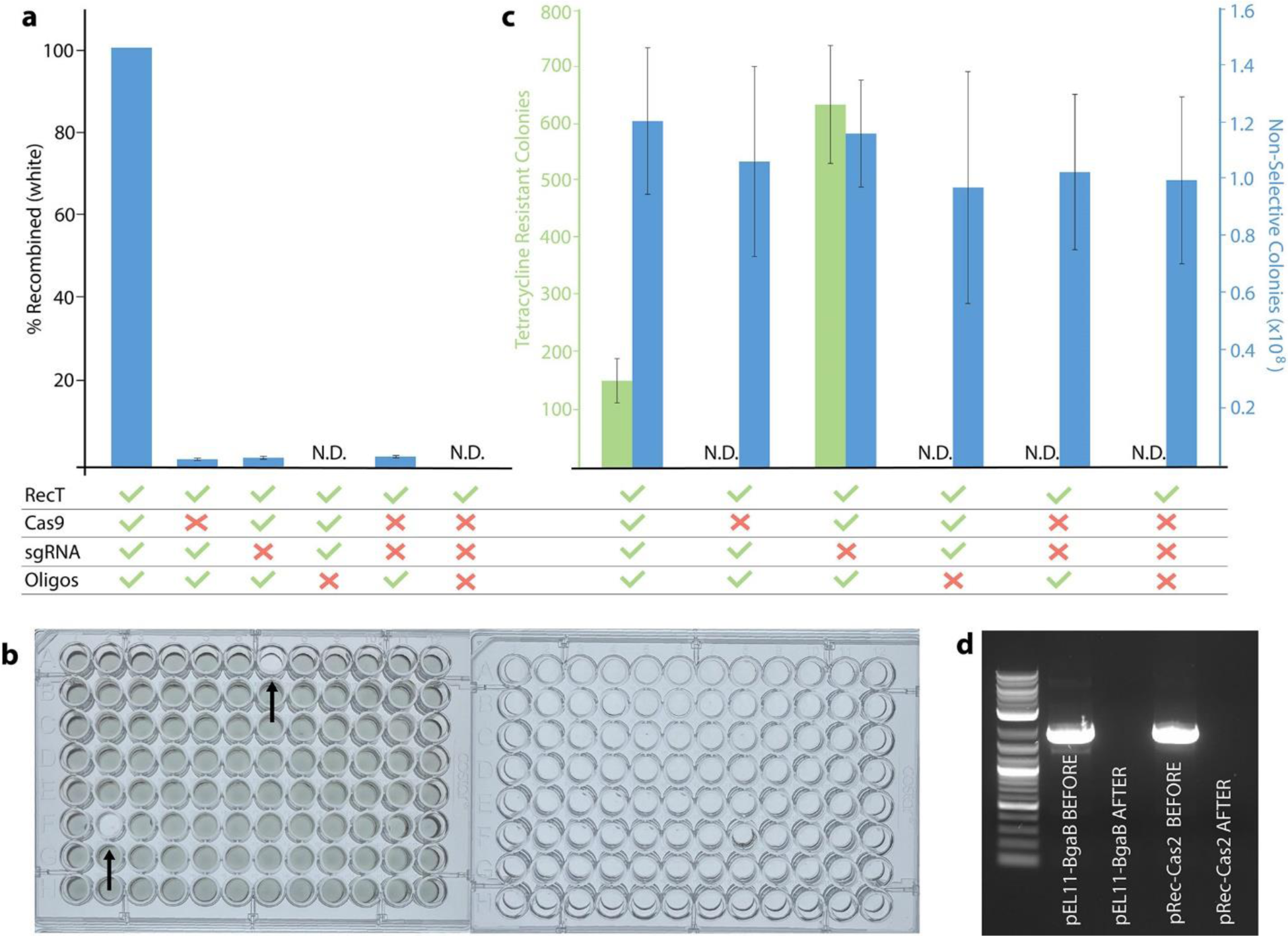
Cas9 counterselection boosts recombineering efficiency to 100%. Error bars represent standard deviation of biological triplicates. a) Recombination efficiency as determined by colony color, b) Representative 96-well plates for colonies picked from condition treated with RecT and recombinogenic oligos but lacking Cas9 and sgRNA (left) and condition with Cas9 and sgRNA (right). Arrows on the left plate indicate recombined strains. Every colony on the right plate lacks BgaB activity. c) Number of colonies obtained on tetracycline and non-selective agar plates showing that on the order of hundreds of recombined colonies may be obtained from a transformation. d) Colony PCR results showing removal of pEL11-BgaB (plasmid encoding RecT and sgRNAs) and pRec-Cas2 (Plasmid encoding Cas9) following counterselection and curing. N.D: not detected.

We desired to eliminate the recombinase plasmid after achieving the targeted mutation, to remove the selection marker and prepare the strain for future genetic engineering work. The additional sgRNA targeting the pEL11 backbone results in Cas9 cleavage of the RecT expression plasmid. Additionally, the Cas9 expression plasmid is maintained by RepH, which has a low copy number in *E. limosum* and is easily lost by curing [13]. Following selection on tetracycline plates, we grew recombined strains for two generations in liquid culture without selection to cure out the Cas9 plasmid, resulting in a final strain encoding the desired genomic point mutation with no plasmids or selection markers (**Figure 4c, Figure S5**). The design of the system makes it easy to swap out guide RNAs on the recombinase plasmid by simply PCR amplifying the full plasmid and encoding a new sgRNA on oligo overhangs. This enables quick turnaround for genome engineering workflows in a strategy which can be applied to any arbitrary target in the genome with a suitable nearby PAM site. RecT recombination with Cas9 counterselection enables rapid and simple generation of genome point mutations in *E. limosum* with up to 100% efficiency, resulting in a final strain containing no plasmids or selection markers.

### Recombinase-Mediated Genome Engineering

As a practical demonstration of our system, we applied the optimized recombineering/Cas9 counterselection scheme to create an *E. limosum* knockout strain incapable of biofilm formation. Extracellular polymeric substance (EPS) in *E. limosum* forms a thick, slimy mass which makes it challenging to manipulate by pipetting and centrifugation and hinders its applications [47]. We previously identified the genes responsible for EPS production and created a knockout strain using RecA-mediated HR, however this process left the *catP* selection marker in the genome, significantly reducing the strain’s usefulness in an engineering context [16]. Here, we generated an analogous strain without any selection markers left in the genome utilizing RecT recombineering with a 120 bp antisense oligo encoding a complete knockout of the operon. We constructed plasmid pEL11-eps encoding RecT and sgRNA targeting the *eps* gene cluster in the genome along with sgRNA for plasmid removal and transformed this into *E. limosum* along with pRec-Cas2 and the recombinogenic oligo in a strategy analogous to that utilized for BgaB inactivation (**Figure 5a**).

**Figure 5:**
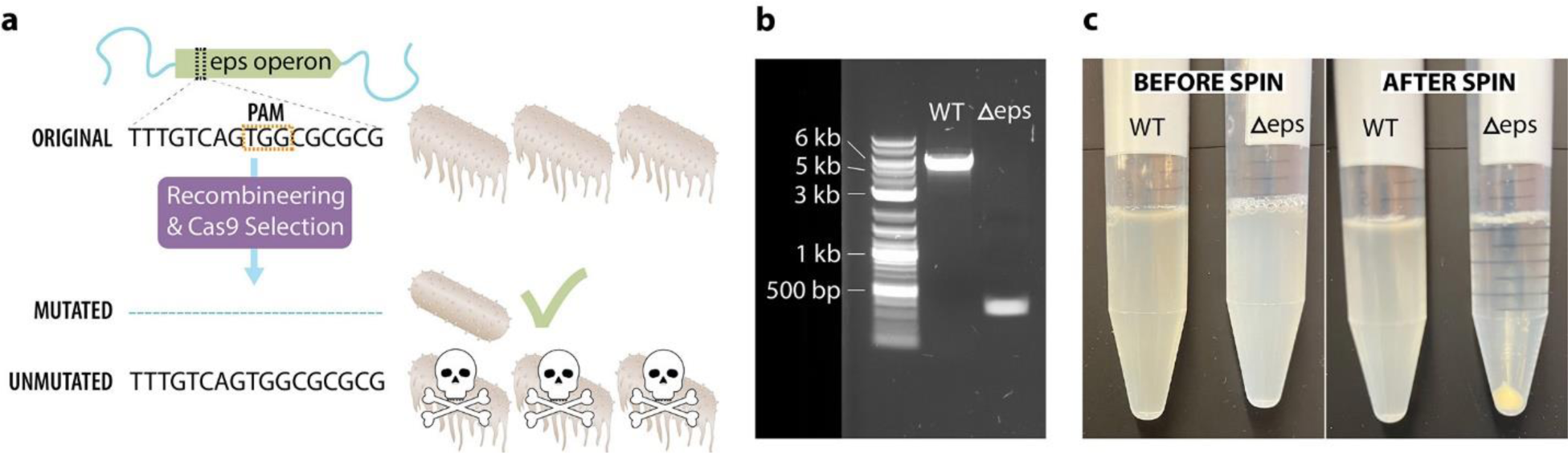
Recombination for biofilm abolition with Cas9 counterselection. a) Scheme for *eps* gene cluster inactivation. b) Colony PCR results for amplification across *eps* gene cluster region +150 bp on either side. Bands show full length gene cluster of ∼5kb for WT strain and removal of operon for engineered strain leaving just the ∼150 bp on either side. c) Centrifugation analysis demonstrating Δ*eps* strain pellets at low speed (3,500 xg). Both strains are fully suspended before centrifugation (left), but only the Δ*eps* strain pellets after centrifugation (right).

Following induction, transformation, and selection, we obtained EPS-deficient strains and performed colony PCR and sequencing to confirm the correct genomic mutation (**Figure 5b**). Since EPS production prevents pelleting by centrifugation at low speed, we utilized this as a readout to assess our knockout strain. As expected, the Δ*eps* strain pelleted fully whereas no pelleting was observed with a wild type culture centrifuged under the same conditions (**Figure 5c**), confirming that we were able to regenerate the Δ*eps* strain using a clean approach leaving no selection markers in the genome.

Rapid genomic modification has been a major bottleneck in the synthetic biology of acetogens, and Clostridia more broadly. The development here of high-efficiency recombineering and markerless genome engineering will significantly streamline the development of engineered *E. limosum* strains for production of an expanded array of products from single-carbon substrates, and facilitate fundamental investigations into acetogen physiology and metabolism. Beyond *E. limosum,* we believe the workflow presented here – specifically the mining of prophages and their original phages for highly active RecT’s – may serve as a roadmap for the broader Clostridial synthetic biology community. Additionally, the recombinase library we generated employs a widely used Clostridial plasmid architecture (pIP404 ori, catP for thiamphenicol resistance, p2tetO1 promoter) and encodes a broad range of RecT’s (including several from Clostridia), facilitating its testing in other Clostridia of industrial significance and importance in human health. Given the wide host range of RecT enzymes, such screening will likely accelerate development of genetic tools for this notoriously recalcitrant class of organisms.

## Materials and Methods

### Reagents, Bacterial Strains, and Growth Conditions

All cloning enzymes were obtained from New England Biolabs (NEB), all oligos 60 bp or shorter were obtained from Genewiz/Azenta and all oligos 61 bp+ or containing phosphorothioate bases were obtained from Sigma-Aldrich. Pre-cloned plasmid constructs utilizing synthesized DNA were obtained from the Joint Genome Institute (JGI). PCR cleanup and miniprep kits were obtained from Zymo Research, gDNA prep kit from NEB, and all other reagents from Fisher Scientific. Cloning and plasmid work were carried out in either *E. coli* NEB5α or stable competent *E. coli*, both from NEB and all work in *E. limosum* was conducted in strain ATCC 8486. *E. coli* was grown at 37°C in liquid LB, or LB agar containing 50 μg/mL ampicillin or chloramphenicol. *E. limosum* was grown anaerobically at 37°C without agitation in DSMZ 135 medium without sulfide for liquid culture and reinforced clostridial medium (RCM) for agar plates. Thiamphenicol was used at a concentration of 5 μg/mL, clarithromycin and tetracycline at 0.5 μg/mL, and aTc at 30 ng/mL. All anaerobic work was done in a Coy Lab Products flexible anaerobic chamber with a 10% CO_2_, 5% H_2_, 85% N_2_ gas mix. Electroporation was conducted using a Bio-Rad GenePulser XCell electroporator following a protocol and electrocompetency preparation reported previously [13]. Optical density and DNA concentration measurements were taken with a Molecular Devices QuickDrop spectrophotometer.

### Cloning and Construct Preparation

Plasmids pEL6-pEL67 were obtained pre-cloned and codon harmonized from JGI. Plasmid pRTS3 was cloned via a 5-piece NEB HiFi assembly using backbone amplified from pUC19 with oligos oPAS222 and 223, overhangs for genomic integration amplified from *E. limosum* ATCC 8486 gDNA using oligos oPAS328 and 271 for the LHS and oPAS269 and 329 for RHS, *Tet(M)* amplified from pVEL4 using oPAS270 and 227, and *ermB’* amplified from synthesized DNA using oPAS267 and 268. Linear DNA for integration into the *E. limosum* genome was amplified from the assembled plasmid using oPAS330 and 331. Control plasmid pRTS3.(+) was constructed in the same way with the exception of using functional *ermB* amplified from pCL2.1 with oPAS267 and 268 rather than mutated *ermB’* [13].

Plasmid pRec-Cas1 was cloned via a 3-piece T4 ligation reaction using backbone amplified from pUC19 with oligos oPAS620 and 621, *RepH* amplified from pMTL83151 using oPAS622 and 623, and *Tet(M)* amplified from pVEL4 using oPAS624 and 625 [48]. Plasmid pRec-Cas2 was then cloned by a 2-piece T4 ligation reaction using Cas9 amplified from synthesized DNA with oPAS698 and 676 and digested with NheI and BsrGI and backbone amplified from pRec-Cas1 with oPA696 and 697 before digest with NheI and BsrGI. pEL11-BgaB was in turn cloned by digesting pEL11 with NheI to linearize and performing an NEB HiFi assembly reaction with a combined sgRNA expression cassette targeting BgaB and pEL11. Individual sgRNA fragments targeting BgaB and pEL11 were synthesized and then combined using overlap extension PCR by amplifying pEL11 sgRNA with oPAS682 and 683 and BgaB sgRNA with 684 and 685 and then combining them and amplifying the double cassette with oPAS717 and 718. Plasmid pEL11-eps was cloned by amplifying pEL11-BgaB with oEPS53 and 54 to replace the BgaB-targeting sgRNA with one targeting the eps operon and then re-ligated in a one-piece NEB-HiFi assembly reaction. Plasmid pBgaBg was cloned via a 2-piece NEB HiFi assembly using a backbone from pRTS3.(+) amplified with oPAS496 and 497, and *bgaB* driven by nifJp amplified from pELIM2.2 using oPAS498 and 499 [13]. Linear DNA for genomic knock-in transformation in *E. limosum* was amplified from the plasmid using oligos oPAS330 and 331.

### Recombinase System Development

*E. limosum* genomes ATCC 8486, SA11, and 81C1 were screening for prophage regions using PHASTER (accession numbers NZ_CP019962.1, NZ_LR215983.1, NZ_CP011914.1 respectively). We selected phages based on gene hit count as determined by PHASTER, assessing ones with four or more clustered gene hits in a single genome region. Phages PLE3, phiCT19406A, phiCT19406B, phiCTC2A, phiCTC2B, and phiCT19406C have accession numbers NC_031125, NC_030950.1, NC_030947.1, NC_030949.1, NC_030951.1, NC_029006.1 respectively. Strain RTS was generated by PCR amplifying linear DNA encoding *ermB’* from pRTS3 and transforming it into WT *E. limosum* utilizing RecA HR for genomic integration as described previously [16]. Recombinase plasmids were transformed into *E. limosum* strain RTS and liquid cultures grown to an OD_600_ of 0.5 before being pooled into a single culture. Any deviations from OD_600_ 0.5 between cultures were normalized to 0.5 with DSMZ 135 media before 750 μL of each were combined into a single culture from which competent cells were prepared. Transforms were performed with 50 μL competent cells and 100 ng/μL DNA with 120 bp oligos in both sense and antisense orientations as well as a no DNA control and plated in a dilution series 1:10, 1:100, 1:1000, 1:10000. Plates were allowed to grow for 6 days before 100 single colonies were picked from each condition. Recombinases encoded by individual resistant colonies were determined by colony PCR to amplify the recombinase gene, and sanger sequencing to determine recombinase identity. Any sequencing samples with a quality score of 24 or below were not included in the dataset. Optimization was performed solely on strain RTS + pEL11 but otherwise in an analogous manner. All transformations were performed in biological triplicate.

### Implementation of Cas9 Counterselection

A homology model of the *Geobacillus stearothermophilus* beta-galactosidase (BgaB) was made with the SWISS-MODEL server, using the crystal structure of the galactose-bound *Bifidobacterium animalis* enzyme (4UNI) as template [49], [50]. Conserved active site residues were identified by overlaying the structures in PYMOL. Strain *E. limosum* pEL11-BgaB/BgaBg was created by performing a knock-in transformation to place *BgaB* with *ermB* for selection into the genome in a fashion and location identical to RTS [16]. Next, competent cells were prepared from this strain and transformed with pEL11-BgaB and selected on thiamphenicol. Finally, competent cells were prepared from this strain with aTc induction and used for recombineering transformations. Recombineering transformations were performed in biological triplicates with 50 μL of competent cells and 10 μL of DNA for a total volume of 60 μL according to **Table 1**. Recombinogenic oligos were used at a final concentration of 1000 ng/μL and plasmid DNA at 4 ng/μL. aTc was added to the recovery media. Colonies were counted after six days of growth and viability determined by comparison to transform B6. 96 colonies were selected from each transformation plate for conditions B1-B6 and grown in 200 μL 96-well plate cultures overnight. Cells were spotted onto tetracycline, thiamphenicol, and non-selective plates and then 250 μg/mL S-Gal and 400 μg/mL ammonium ferric citrate added to each well in the 96-well culture plates and allowed to incubate at 37°C for 1 hour before visually assessing BgaB activity. After correctly recombined strains were identified, they were subcultured twice in non-selective liquid media before plating and colony PCR analysis to confirm loss of pEL11-BgaB and pRec-Cas2.

**Table 1:** Transformations for Cas9 counterselection demonstration using the BgaB reporter.

| Transform | Strain | DNA | Plating |
| --- | --- | --- | --- |
| B1 | pEL11/BgaBg | oPAS518 | RCM + tetracycline & RCM only |
| B2 | pEL11-BgaB/BgaBg | oPAS518<br>pRec-Cas2 | RCM + tetracycline & RCM only |
| B3 | pEL11/BgaBg | oPAS518<br>pRec-Cas2 | RCM + tetracycline & RCM only |
| B4 | pEL11-BgaB/BgaBg | pRec-Cas2 | RCM + tetracycline & RCM only |
| B5 | pEL11-BgaB/BgaBg | oPAS518 | RCM + tetracycline & RCM only |
| B6 | pEL11/BgaBg | None | RCM + tetracycline & RCM only |

### Recombinase-mediated Genome Engineering

Wild type competent cells were transformed with pEL11-eps, selected, cultured, induced, and made competent. 50μL pEL11-eps competent cells were transformed with 5μL oligo oEPS55 at a final concentration of 500 ng/μL and plasmid pRec-Cas2 at 4 ng/μL with aTc added to the recovery media for cas9 expression before plating on tetracycline plates. After six days of growth, single colonies were picked from agar plates into a well plate with 150μL DSMZ 135 media per well and allowed to grow overnight. Following growth, each well was pipetted to qualitatively assess EPS production, and cells from a well without any biofilm were passaged into a 5mL culture and allowed to grow overnight. Cultures were then centrifuged at 3500xg for 5 minutes to assess pelleting. Colony PCR was performed on the knockout region using oligos oPAS763 and 764.

## Supporting information

All supplemental information

## Author Information

### Author Contributions

PAS devised and performed experiments and engineering strategies, wrote the manuscript, and created the figures and graphics. Both authors analyzed data. BMW conceived of the concept and edited the manuscript and figures.

## Acknowledgements

This work was financially supported by Northeastern University startup funds and Department of Energy, Advanced Research Projects Agency-Energy ECOSynBIO under award number #DE-AR0001511. The work (proposal: https://doi.org/10.46936/10.25585/60008422) conducted by the U.S. Department of Energy Joint Genome Institute (https://ror.org/04xm1d337), a DOE Office of Science User Facility, is supported by the Office of Science of the U.S. Department of Energy operated under Contract No. DE-AC02-05CH11231.

## References

[1] A. Mukherjee, C. Lordan, R. P. Ross, and P. D. Cotter, “Gut microbes from the phylogenetically diverse genus Eubacterium and their various contributions to gut health,” Gut Microbes, vol. 12, no. 1, 2020, doi: 10.1080/19490976.2020.1802866.

[2] J. Y. Kim et al., “Methanol supply speeds up synthesis gas fermentation by methylotrophic-acetogenic bacterium, Eubacterium limosum KIST612,” Bioresour. Technol., vol. 321, no. November 2020, p. 124521, 2021, doi: 10.1016/j.biortech.2020.124521.

[3] S. Jin et al., “Synthetic biology on acetogenic bacteria for highly efficient conversion of c1 gases to biochemicals,” Int. J. Mol. Sci., vol. 21, no. 20, pp. 1–25, 2020, doi: 10.3390/ijms21207639.

[4] T. Ueki, K. P. Nevin, T. L. Woodard, and D. R. Lovley, “Converting carbon dioxide to butyrate with an engineered strain of Clostridium ljungdahlii,” MBio, vol. 5, no. 5, pp. 19–23, 2014, doi: 10.1128/mBio.01636-14.

[5] M. Flaiz, G. Ludwig, F. R. Bengelsdorf, and P. Dürre, “Production of the biocommodities butanol and acetone from methanol with fluorescent FAST-tagged proteins using metabolically engineered strains of Eubacterium limosum,” Biotechnol. Biofuels, vol. 14, no. 1, pp. 1–20, 2021, doi: 10.1186/s13068-021-01966-2.

[6] B. Karlson, C. Bellavitis, and N. France, “Commercializing LanzaTech, from waste to fuel: An effectuation case,” J. Manag. Organ., vol. 27, no. 1, pp. 175–196, 2021, doi: 10.1017/jmo.2017.83.

[7] C. A. Cotton, N. J. Claassens, S. Benito-Vaquerizo, and A. Bar-Even, “Renewable methanol and formate as microbial feedstocks,” Curr. Opin. Biotechnol., vol. 62, pp. 168–180, 2020, doi: 10.1016/j.copbio.2019.10.002.

[8] A. Braune and M. Blaut, “Bacterial species involved in the conversion of dietary flavonoids in the human gut,” Gut Microbes, vol. 7, no. 3, pp. 216–234, 2016.

[9] J. B. Ellenbogen, R. Jiang, D. J. Kountz, L. Zhang, and J. A. Krzycki, “The MttB superfamily member MtyB from the human gut symbiont Eubacterium limosum is a cobalamin-dependent γ-butyrobetaine methyltransferase,” J. Biol. Chem., vol. 297, no. 5, p. 101327, 2021, doi: 10.1016/j.jbc.2021.101327.

[10] J. Shin et al., “Genome Engineering of Eubacterium limosum Using Expanded Genetic Tools and the CRISPR-Cas9 System,” ACS Synth. Biol., vol. 8, no. 9, pp. 2059–2068, 2019, doi: 10.1021/acssynbio.9b00150.

[11] Y. Song et al., “Determination of the Genome and Primary Transcriptome of Syngas Fermenting Eubacterium limosum ATCC,” Sci. Rep., vol. 7, no. 1, pp. 1–11, 2017, doi: 10.1038/s41598-017-14123-3.

[12] Y. Song et al., “Development of highly characterized genetic bioparts for efficient gene expression in CO 2 -fixing Eubacterium limosum,” Metab. Eng., vol. 72, no. March, pp. 215–226, 2022, doi: 10.1016/j.ymben.2022.03.016.

[13] P. A. Sanford and B. M. Woolston, “Expanding the genetic engineering toolbox for the metabolically flexible acetogen Eubacterium limosum,” J. Ind. Microbiol. Biotechnol., no. July, 2022, doi: 10.1093/jimb/kuac019.

[14] J. Shin et al., “Genome-wide CRISPRi screen identifies enhanced autolithotrophic phenotypes in acetogenic bacterium Eubacterium limosum,” PNAS, vol. 120, no. 6, 2023.

[15] J. Millard, A. Agius, Y. Zhang, P. Soucaille, and N. P. Minton, “Exploitation of a Type 1 Toxin–Antitoxin System as an Inducible Counter-Selective Marker for Genome Editing in the Acetogen Eubacterium limosum,” Microorganisms, vol. 11, no. 5, pp. 1–13, 2023, doi: 10.3390/microorganisms11051256.

[16] P. A. Sanford, K. G. Miller, K. O. Hoyt, and B. M. Woolston, “Deletion of biofilm synthesis in Eubacterium limosum ATCC 8486 improves handling and transformation efficiency,” FEMS Microbiol. Lett., vol. 370, pp. 1–8, 2023, doi: 10.1093/femsle/fnad030.

[17] C. A. Vees, C. S. Neuendorf, and S. Pflügl, “Towards continuous industrial bioprocessing with solventogenic and acetogenic clostridia: challenges, progress and perspectives,” J. Ind. Microbiol. Biotechnol., vol. 47, no. 9–10, pp. 753–787, 2020, doi: 10.1007/s10295-020-02296-2.

[18] K. Charubin, R. K. Bennett, A. G. Fast, and E. T. Papoutsakis, “Engineering Clostridium organisms as microbial cell-factories: challenges & opportunities,” Metab. Eng., vol. 50, no. June, pp. 173–191, 2018, doi: 10.1016/j.ymben.2018.07.012.

[19] P. A. Sanford and B. M. Woolston, “Synthetic or natural? Metabolic engineering for assimilation and valorization of methanol,” Curr. Opin. Biotechnol., vol. 74, pp. 171–179, 2022, doi: 10.1016/j.copbio.2021.12.001.

[20] G. Pines, E. F. Freed, J. D. Winkler, and R. T. Gill, “Bacterial Recombineering: Genome Engineering via Phage-Based Homologous Recombination,” ACS Synth. Biol., vol. 4, no. 11, pp. 1176–1185, 2015, doi: 10.1021/acssynbio.5b00009.

[21] K. C. Murphy and K. G. Campellone, “Lambda Red-mediated recombinogenic engineering of enterohemorrhagic and enteropathogenic E. coli,” BMC Mol. Biol., vol. 4, pp. 1–12, 2003, doi: 10.1186/1471-2199-4-11.

[22] H. H. Wang et al., “Programming cells by multiplex genome engineering and accelerated evolution,” Nature, vol. 460, no. 7257, pp. 894–898, 2009, doi: 10.1038/nature08187.

[23] R. Kolodner, S. D. Hall, and C. Luisi-DeLuca, “Homologous pairing proteins encoded by the Escherichia coii recE and recT genes,” Mol. Microbiol., vol. 11, no. 1, pp. 23–30, 1994, doi: 10.1111/j.1365-2958.1994.tb00286.x.

[24] M. E. Pyne, M. Moo-Young, D. A. Chung, and C. P. Chou, “Coupling the CRISPR/Cas9 system with lambda red recombineering enables simplified chromosomal gene replacement in Escherichia coli,” Appl. Environ. Microbiol., vol. 81, no. 15, pp. 5103–5114, 2015, doi: 10.1128/AEM.01248-15.

[25] C. R. Reisch and K. L. J. Prather, “The no-SCAR (Scarless Cas9 Assisted Recombineering) system for genome editing in Escherichia coli,” Sci. Rep., vol. 5, no. October, pp. 1–12, 2015, doi: 10.1038/srep15096.

[26] H. M. Ellis, D. Yu, T. DiTizio, and D. L. Court, “High efficiency mutagenesis, repair, and engineering of chromosomal DNA using single-stranded oligonucleotides,” Proc. Natl. Acad. Sci. U. S. A., vol. 98, no. 12, pp. 6742–6746, 2001, doi: 10.1073/pnas.121164898.

[27] T. M. Wannier et al., “Improved bacterial recombineering by parallelized protein discovery,” Proc. Natl. Acad. Sci. U. S. A., vol. 117, no. 24, pp. 13689–13698, 2020, doi: 10.1073/pnas.2001588117.

[28] S. Datta, N. Costantino, X. Zhou, and D. L. Court, “Identification and analysis of recombineering functions from Gram-negative and Gram-positive bacteria and their phages,” Proc. Natl. Acad. Sci. U. S. A., vol. 105, no. 5, pp. 1626–1631, 2008, doi: 10.1073/pnas.0709089105.

[29] P. Noirot and R. D. Kolodner, “DNA strand invasion promoted by Escherichia coli RecT protein,” J. Biol. Chem., vol. 273, no. 20, pp. 12274–12280, 1998, doi: 10.1074/jbc.273.20.12274.

[30] A. Roca, M. Cox, and S. Brenner, “The RecA Protein: Structure and Function,” Crit. Rev. Biochem. Mol. Biol., vol. 25, no. 6, pp. 415–456, 1990, doi: 10.3109/10409239009090617.

[31] H. Dong, W. Tao, F. Gong, Y. Li, and Y. Zhang, “A functional recT gene for recombineering of Clostridium,” J. Biotechnol., vol. 173, no. 1, pp. 65–67, 2014, doi: 10.1016/j.jbiotec.2013.12.011.

[32] B. Swingle, Z. Bao, E. Markel, A. Chambers, and S. Cartinhour, “Recombineering using recTE from pseudomonas syringae,” Appl. Environ. Microbiol., vol. 76, no. 15, pp. 4960–4968, 2010, doi: 10.1128/AEM.00911-10.

[33] D. Arndt et al., “PHASTER: a better, faster version of the PHAST phage search tool,” Nucleic Acids Res., vol. 44, no. W1, pp. W16–W21, 2016, doi: 10.1093/nar/gkw387.

[34] C. Canchaya, C. Proux, G. Fournous, A. Bruttin, and H. Brüssow, “Prophage Genomics,” Microbiol. Mol. Biol. Rev., vol. 67, no. 3, pp. 473–473, 2003, doi: 10.1128/mmbr.67.3.473.2003.

[35] H. Dong, W. Tao, Y. Zhang, and Y. Li, “Development of an anhydrotetracycline-inducible gene expression system for solvent-producing Clostridium acetobutylicum: A useful tool for strain engineering,” Metab. Eng., vol. 14, no. 1, pp. 59–67, 2012, doi: 10.1016/j.ymben.2011.10.004.

[36] T. Aparicio, V. de Lorenzo, and E. Martínez-García, “CRISPR/Cas9-enhanced ssDNA recombineering for Pseudomonas putida,” Microb. Biotechnol., vol. 12, no. 5, pp. 1076–1089, 2019, doi: 10.1111/1751-7915.13453.

[37] C. Wyman, D. Ristic, and R. Kanaar, “Homologous recombination-mediated double-strand break repair,” DNA Repair (Amst)., vol. 3, no. 8–9, pp. 827–833, 2004, doi: 10.1016/j.dnarep.2004.03.037.

[38] J. A. Sawitzke et al., “Probing cellular processes with oligo-mediated recombination and using the knowledge gained to optimize recombineering,” J. Mol. Biol., vol. 407, no. 1, pp. 45–59, 2011, doi: 10.1016/j.jmb.2011.01.030.

[39] M. Nakayama and O. Ohara, “Improvement of Recombination Efficiency by Mutation of Red Proteins,” Biotechniques, vol. 38, no. 6, pp. 917–924, 2005, doi: 10.2144/05386RR02.

[40] Y. Ding et al., “Increasing the homologous recombination efficiency of eukaryotic microorganisms for enhanced genome engineering,” Appl. Microbiol. Biotechnol., vol. 103, no. 11, pp. 4313–4324, 2019, doi: 10.1007/s00253-019-09802-2.

[41] J. Ren, S. Karna, H. M. Lee, S. M. Yoo, and D. Na, “Artificial transformation methodologies for improving the efficiency of plasmid DNA transformation and simplifying its use,” Appl. Microbiol. Biotechnol., vol. 103, no. 23–24, pp. 9205–9215, 2019, doi: 10.1007/s00253-019-10173-x.

[42] B. E. Dutra, V. A. Sutera, and S. T. Lovett, “RecA-independent recombination is efficient but limited by exonucleases,” Proc. Natl. Acad. Sci. U. S. A., vol. 104, no. 1, pp. 216–221, 2007, doi: 10.1073/pnas.0608293104.

[43] S. Spitzer and F. Eckstein, “Inhibition of deoxyribonucleases by phosphorothioate groups in oligodeoxyribonucleotides,” Nucleic Acids Res., vol. 16, no. 24, pp. 11691–11704, 1988.

[44] Y. Jiang, B. Chen, C. Duan, B. Sun, J. Yang, and S. Yang, “Multigene editing in the Escherichia coli genome via the CRISPR-Cas9 system,” Appl. Environ. Microbiol., vol. 81, no. 7, pp. 2506–2514, 2015, doi: 10.1128/AEM.04023-14.

[45] J. Yuan, M. Martinez-Bilbao, and R. E. Huber, “Substitutions for Glu-537 of β-galactosidase from Escherichia coli cause large decreases in catalytic activity,” Biochem. J., vol. 299, no. 2, pp. 527–531, 1994, doi: 10.1042/bj2990527.

[46] K. L. Morrison and G. A. Weiss, “Combinatorial alanine-scanning,” Curr. Opin. Chem. Biol., vol. 5, no. 3, pp. 302–307, 2001, doi: 10.1016/S1367-5931(00)00206-4.

[47] P. Loubiere, E. Gros, V. Paquet, and N. D. Lindley, “Kinetics and physiological implications of the growth behaviour of Eubacterium limosum on glucose/methanol mixtures,” J. Gen. Microbiol., vol. 138, no. 5, pp. 979–985, 1992, doi: 10.1099/00221287-138-5-979.

[48] J. T. Heap, O. J. Pennington, S. T. Cartman, and N. P. Minton, “A modular system for Clostridium shuttle plasmids,” J. Microbiol. Methods, vol. 78, no. 1, pp. 79–85, 2009, doi: 10.1016/j.mimet.2009.05.004.

[49] A. Waterhouse et al., “SWISS-MODEL: Homology modelling of protein structures and complexes,” Nucleic Acids Res., vol. 46, no. W1, pp. W296–W303, 2018, doi: 10.1093/nar/gky427.

[50] A. H. Viborg et al., “A β1-6/β1-3 galactosidase from Bifidobacterium animalis subsp. lactisBl-04 gives insight into sub-specificities of β-galactoside catabolism within Bifidobacterium,” Mol. Microbiol., vol. 94, no. 5, pp. 1024–1040, 2014, doi: 10.1111/mmi.12815.

