## Supplementary material for "Development of a recombineering system for the acetogen *Eubacterium limosum* with Cas9 counterselection for markerless genome engineering": All supplemental information

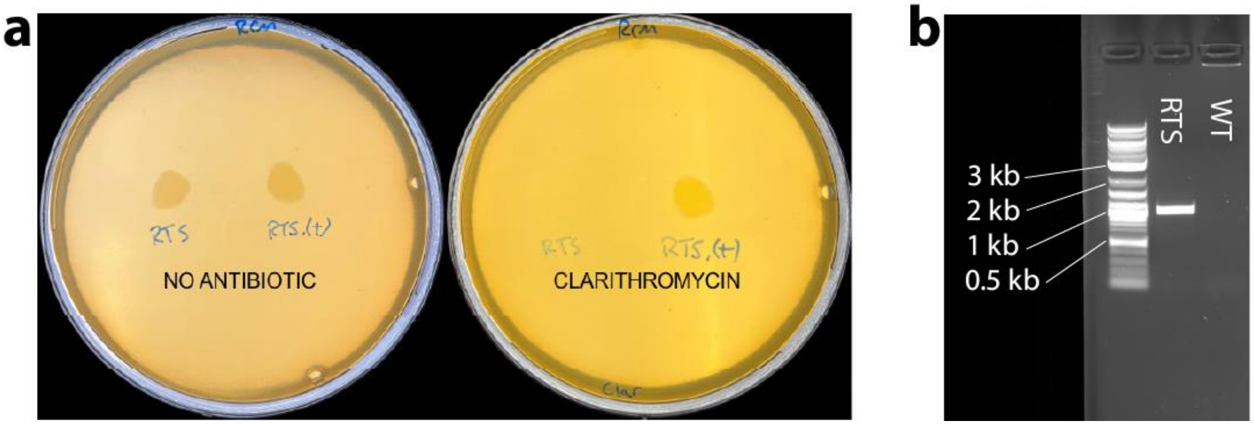

**Figure S1: Creation and confirmation of recombinase test strain.** a) Colony PCR confirmation of *ermB'* genomic integration. PCR from within *ermB'* across junction into genome shows band for RTS but not for WT. b) Growth of strains RTS and RTS.(+) on RCM agar plates with and without clarithromycin. RTS contains the mutated *ermB'* which possesses stop codons early in the coding sequence to prevent expression. Strain RTS.(+) encodes functional *ermB*. Note how RTS is incapable of growth on clarithromycin but RTS.(+) forms a regular spot from liquid culture.

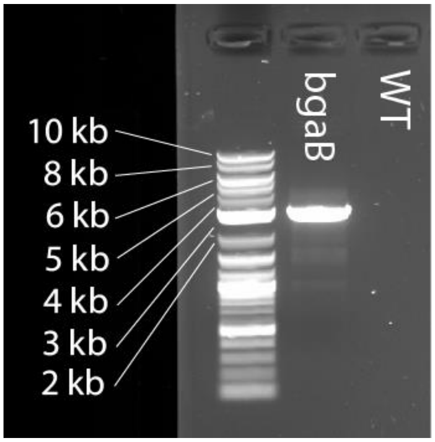

**Figure S2:** Colony PCR confirmation of *bgaB* genomic integration. PCR from within *bgaB* across junction into genome shows band for BgaBg strain but not for WT.

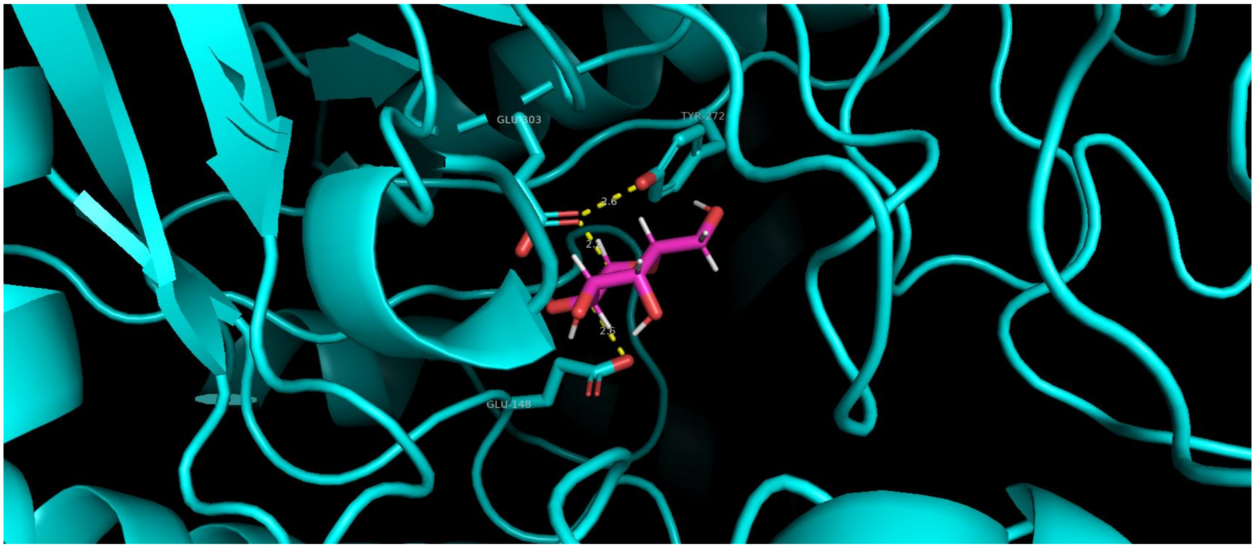

**Figure S3: Identification of catalytic nucleophile by homology modeling**. Homology model of *G. stearothermmophilus* (blue) with galactose docked in active site. Y272, E303, and E148 form the canonical catalytic triad of glycoside hydrolases.

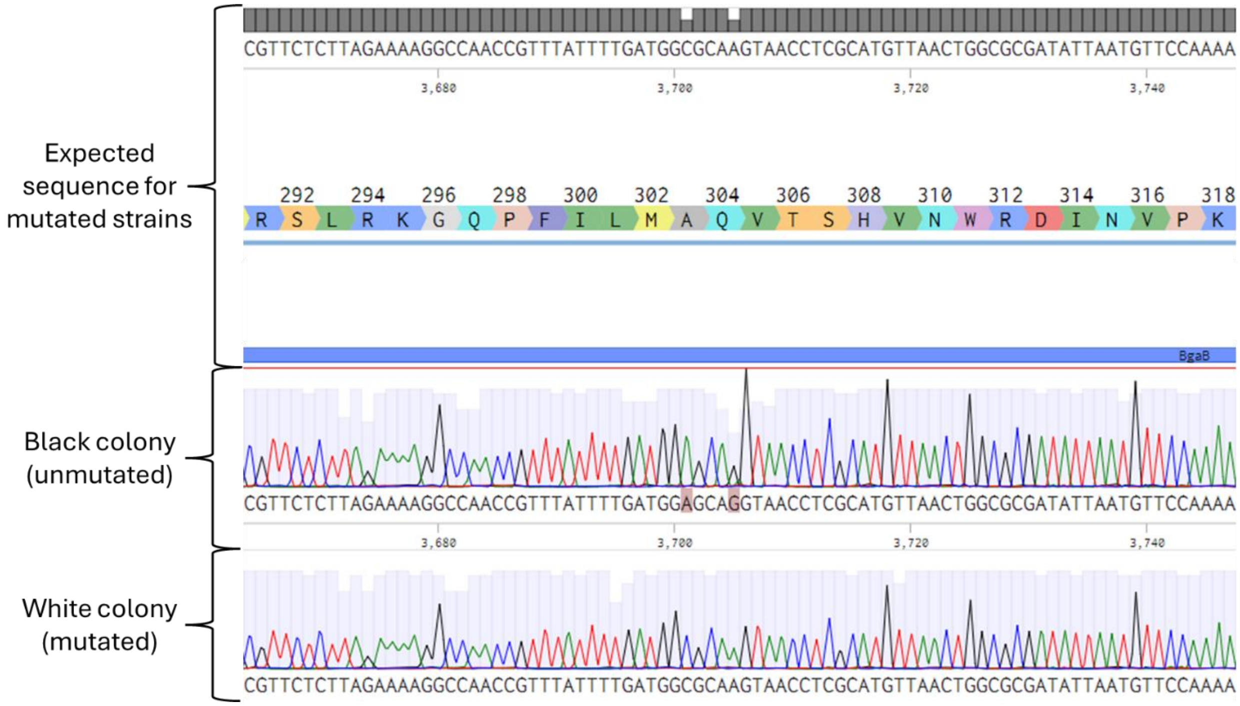

**Figure S4:** Sequencing alignment confirming BgaB genomic mutation and PAM site removal.

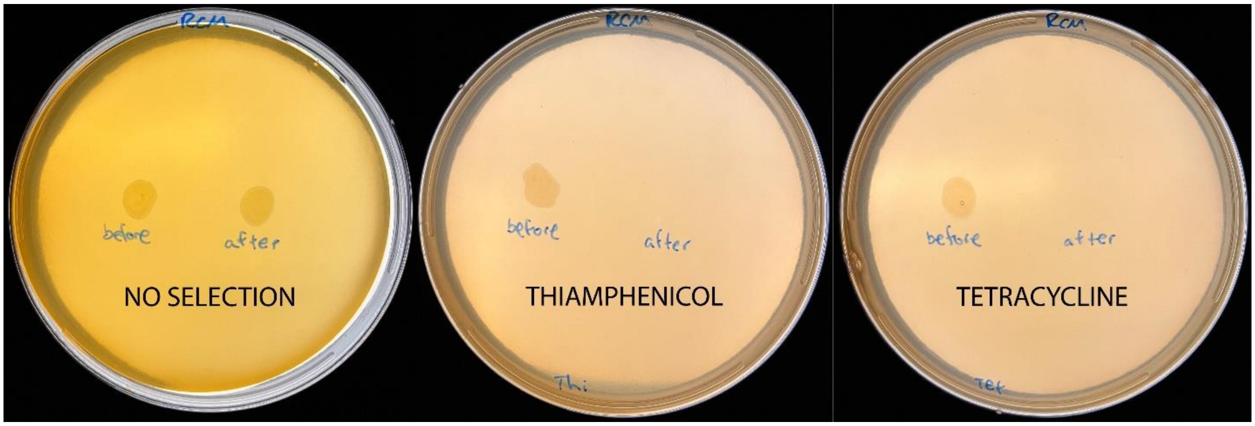

**Figure S5:** Plate spotting assay showing loss of pEL11-BgaB and pRec-Cas2 following counterselection and curing.

**Table S1:** Recombinase list

| **Phylogenetic Source** | **Name** | **Activity** | **Construct** |
|---|---|---|---|
| Clostridium phage phiCT19406A | RecT family recombinase | Putative | pEL6 |
| Clostridium phage phiCT19406B | RecT family recombinase | Putative | pEL7 |
| Clostridium phage phiCTC2A | RecT family recombinase | Putative | pEL8 |
| Clostridium phage phiCTC2B | RecT family recombinase | Putative | pEL9 |
| Clostridium phage phiCT19406C | hypothetical protein | Putative | pEL10 |
| Eubacterium limosum (SA11) | RecT | Putative | pEL11 |
| Eubacterium limosum (81C1) | RecT | Putative | pEL12 |
| Clostridium paraputrificum | recombinase RecT | Putative | pEL13 |
| Clostridium tertium | recombinase RecT | Putative | pEL14 |
| Clostridium carnis | recombinase RecT | Putative | pEL15 |
| Clostridium cuniculi | recombinase RecT | Putative | pEL16 |
| Clostridiales bacterium | recombinase RecT | Putative | pEL17 |
| Clostridium thermopalmarium | recombinase RecT | Putative | pEL18 |
| Caloranaerobacter azorensis | recombinase RecT | Putative | pEL19 |
| Bacillus pseudomycoides | recombinase RecT | Putative | pEL20 |

| | | | |
|---|---|---|---|
| Ureibacillus thermophilus | recombinase RecT | Putative | pEL21 |
| Geobacillus icigianus | recombinase RecT | Putative | pEL23 |
| Brevibacillus halotolerans | recombinase RecT | Putative | pEL24 |
| Listeria booriae | recombinase RecT | Putative | pEL25 |
| Listeria monocytogenes | recombinase RecT | Putative | pEL26 |
| Listeria rocourtiae | recombinase RecT | Putative | pEL27 |
| Paenibacillus alvei | recombinase RecT | Putative | pEL28 |
| Priestia abyssalis | recombinase RecT | Putative | pEL29 |
| Sporolactobacillus terrae | recombinase RecT | Putative | pEL30 |
| Oceanobacillus kimchii | recombinase RecT | Putative | pEL31 |
| Bacillus aquimaris | recombinase RecT | Putative | pEL32 |
| Bacillus haikouensis | recombinase RecT | Putative | pEL33 |
| Priestia megaterium | recombinase RecT | Putative | pEL34 |
| Indiicoccus explosivorum | recombinase RecT | Putative | pEL35 |
| Bacillus enclensis | recombinase RecT | Putative | pEL36 |
| Bacillus velezensis | recombinase RecT | Putative | pEL37 |
| Bacillus spizizenii | recombinase RecT | Putative | pEL38 |
| Bacillus halotolerans | recombinase RecT | Putative | pEL39 |
| Bacillus circulans | recombinase RecT | Putative | pEL40 |
| Bacillus taxi | recombinase RecT | Putative | pEL42 |
| Weizmannia coagulans | recombinase RecT | Putative | pEL44 |
| Escherichia phage TL-2011c | RecT | Putative | pEL45 |
| Exiguobacterium phage vB_Eau M-23 | RecT | Putative | pEL46 |
| Staphylococcus phage 23MRA | RecT | Putative | pEL47 |
| Salmonella phage epsilon15 | RecT | Putative | pEL48 |
| Geobacillus phage GBK2 | RecT | Putative | pEL49 |

| | | | |
|---|---|---|---|
| Mycobacterium phage Phayonce | RecT | Putative | pEL50 |
| Klebsiella phage 5 LV-2017 | RecT | Putative | pEL51 |
| Lactococcus phage bIL309 | RecT | Putative | pEL52 |
| Cronobacter phage PhiCS01 | RecT | Putative | pEL53 |
| Hathewaya histolytica | RecT | Putative | pEL55 |
| Clostridium sporogenes | RecT | Putative | pEL56 |
| Pseudomonas fluorescens R124 | RecT | Putative | pEL57 |
| Proteus mirabilis | recombination protein RecT | Putative | pEL59 |
| Aquitalea aquatilis | recombination protein RecT | Putative | pEL60 |
| Sneathia vaginalis | recombination protein RecT | Putative | pEL61 |
| Escherichia phage Rac-SA53 | recT | Putative | pEL62 |
| Clostridium perfringnes | RecT | Reported (H. Dong, W. Tao, F. Gong, Y. Li, and Y. Zhang, “A functional recT gene for recombineering of Clostridium,” J. Biotechnol., vol. 173, no. 1, pp. 65–67, 2014, doi: 10.1016/j.jbiotec.2013.12.011.) | pEL63 |
| Pseudomonas syringae | RecT | Reported (B. Swingle, Z. Bao, E. Markel, A. Chambers, and S. Cartinhour, “Recombineering using recTE from pseudomonas syringae,” Appl. Environ. Microbiol., vol. 76, no. 15, pp. 4960–4968, 2010, doi: 10.1128/AEM.00911-10.) | pEL64 |
| Bacillus subtilis | GP35 | Reported (S. Datta, N. Costantino, X. Zhou, and D. L. Court, “Identification and analysis of recombineering functions from Gram-negative and Gram-positive bacteria and their phages,” Proc. Natl. Acad. Sci. U. S. A., vol. 105, no. 5, pp. 1626–1631, 2008, doi: 10.1073/pnas.0709089105.) | pEL65 |
| Escherichia coli | RecT | Reported (T. M. Wannier et al., “Improved bacterial | pEL66 |

| | | | |
|---|---|---|---|
| | | recombineering by parallelized protein discovery," Proc. Natl. Acad. Sci. U. S. A., vol. 117, no. 24, pp. 13689–13698, 2020, doi: 10.1073/pnas.2001588117.) | |
| Escherichia coli | Beta | Reported (S. Datta, N. Costantino, X. Zhou, and D. L. Court, "Identification and analysis of recombineering functions from Gram-negative and Gram-positive bacteria and their phages," Proc. Natl. Acad. Sci. U. S. A., vol. 105, no. 5, pp. 1626–1631, 2008, doi: 10.1073/pnas.0709089105.) | pEL67 |

**Table S2:** Plasmids used in this study

| **Construct** | **Description** | **Source** |
|---|---|---|
| pEL6 | recombinase plasmid | This study |
| pEL7 | recombinase plasmid | This study |
| pEL8 | recombinase plasmid | This study |
| pEL9 | recombinase plasmid | This study |
| pEL10 | recombinase plasmid | This study |
| pEL11 | recombinase plasmid | This study |
| pEL12 | recombinase plasmid | This study |
| pEL13 | recombinase plasmid | This study |
| pEL14 | recombinase plasmid | This study |
| pEL15 | recombinase plasmid | This study |
| pEL16 | recombinase plasmid | This study |
| pEL17 | recombinase plasmid | This study |
| pEL18 | recombinase plasmid | This study |
| pEL19 | recombinase plasmid | This study |
| pEL20 | recombinase plasmid | This study |
| pEL21 | recombinase plasmid | This study |
| pEL23 | recombinase plasmid | This study |
| pEL24 | recombinase plasmid | This study |
| pEL25 | recombinase plasmid | This study |
| pEL26 | recombinase plasmid | This study |
| pEL27 | recombinase plasmid | This study |
| pEL28 | recombinase plasmid | This study |

| | | |
|---|---|---|
| pEL29 | recombinase plasmid | This study |
| pEL30 | recombinase plasmid | This study |
| pEL31 | recombinase plasmid | This study |
| pEL32 | recombinase plasmid | This study |
| pEL33 | recombinase plasmid | This study |
| pEL34 | recombinase plasmid | This study |
| pEL35 | recombinase plasmid | This study |
| pEL36 | recombinase plasmid | This study |
| pEL37 | recombinase plasmid | This study |
| pEL38 | recombinase plasmid | This study |
| pEL39 | recombinase plasmid | This study |
| pEL40 | recombinase plasmid | This study |
| pEL42 | recombinase plasmid | This study |
| pEL44 | recombinase plasmid | This study |
| pEL45 | recombinase plasmid | This study |
| pEL46 | recombinase plasmid | This study |
| pEL47 | recombinase plasmid | This study |
| pEL48 | recombinase plasmid | This study |
| pEL49 | recombinase plasmid | This study |
| pEL50 | recombinase plasmid | This study |
| pEL51 | recombinase plasmid | This study |
| pEL52 | recombinase plasmid | This study |
| pEL53 | recombinase plasmid | This study |
| pEL55 | recombinase plasmid | This study |
| pEL56 | recombinase plasmid | This study |
| pEL57 | recombinase plasmid | This study |
| pEL59 | recombinase plasmid | This study |
| pEL60 | recombinase plasmid | This study |
| pEL61 | recombinase plasmid | This study |
| pEL62 | recombinase plasmid | This study |
| pEL63 | recombinase plasmid | This study |
| pEL64 | recombinase plasmid | This study |
| pEL65 | recombinase plasmid | This study |
| pEL66 | recombinase plasmid | This study |
| pEL67 | recombinase plasmid | This study |
| pRec-Cas1 | base plasmid for sequential cloning of Cas9 selection plasmids encoding ColE1 and ampR for E. coli, and Tet(M) and repH for E. limosum | This study |
| pRec-Cas2 | Added Cas9 to pRec-Cas1 under the control of p2TetO1 | This study |
| pBgaBg | Plasmid encoding knock-in DNA to insert *BgaB* into E. limosm genome | This study |

| | | |
|---|---|---|
| pEL11-BgaB | Cas9 counterselection plasmid for *BgaB* mutation, added two sgRNAs to pEL11 | This study |
| pEL2.1 | Expression of CatP in *E. limosum* under the control of nifJp | Sanford & Woolston "Expanding the genetic engineering toolbox for the metabolically flexible acetogen Eubacterium limosum ," J. Ind. Microbiol. Biotechnol., no. July, 2022, doi: 10.1093/jimb/kuac019. |
| pEL3.1 | Expression of CatP in *E. limosum* under the control of p2TetO1 | Sanford & Woolston "Expanding the genetic engineering toolbox for the metabolically flexible acetogen Eubacterium limosum ," J. Ind. Microbiol. Biotechnol., no. July, 2022, doi: 10.1093/jimb/kuac019. |
| pMTL82254 | Backbone for cloning Cas9 counterselection constructs | Heap, J. T., Pennington, O. J., Cartman, S. T., & Minton, N. P. "A modular system for Clostridium shuttle plasmids" Journal of Microbiological Methods, 78(1), 79–85. 2009 https://doi.org/10.1016/j.mimet.2009.05.004 |
| pRTS3 | Plasmid encoding knock-in DNA encoding *ermB'* for recombinase library screening | This study |
| pRTS3.(+) | Positive control for RTS3 with a functional *ermB* gene. | This study |
| pUC19 | Backbone for cloning of other constructs | NEB |
| pVEL4 | Source of *Tet(M)* for cloning of other constructs | Sanford & Woolston "Expanding the genetic engineering toolbox for the metabolically flexible acetogen Eubacterium limosum ," J. Ind. Microbiol. Biotechnol., no. July, 2022, doi: 10.1093/jimb/kuac019. |
| pCL2.1 | Source of *ermB* for cloning pRTS3.(+) | Sanford & Woolston "Expanding the genetic engineering toolbox for the metabolically flexible acetogen Eubacterium limosum ," J. Ind. Microbiol. Biotechnol., no. July, 2022, doi: 10.1093/jimb/kuac019. |
| pELIM2.2 | Source of *BgaB* for cloning pBgaBg | Sanford & Woolston "Expanding the genetic engineering toolbox for the metabolically flexible acetogen Eubacterium limosum ," J. Ind. Microbiol. Biotechnol., no. July, 2022, doi: 10.1093/jimb/kuac019. |
| pEL11-eps | Cas9 counterselection plasmid for eps operon inactivation. Identical to | This study |

| | |
|---|---|
| | pEL11-BgaB other than the target sgRNA |

**Table S3:** Oligos used in this study

| Name | Sequence | Description |
|---|---|---|
| oPAS222 | GTATTGGGCGCTCTTCCG | Amplify pUC19 backbone for pRTS3 and pRTS3.(+) |
| oPAS223 | GACGAAAGGGCCTCGTGA | Amplify pUC19 backbone for pRTS3 and pRTS3.(+) |
| oPAS328 | AGGCGTATCACGAGGCCCTTTCGTCcattaataagagagaggtactcattatgaattaca | Amplify overhangs for knock-in of pRTS3/pRTS3.(+) |
| oPAS271 | aaggatctcaacggcctatttggcctaaatgaaaccatattcgaaaaaatcgtctgtctg | Amplify overhangs for knock-in of pRTS3/pRTS3.(+) |
| oPAS269 | ttacacgttactaaagggaatgtgtttaatgttcaggccataaaattaagcgc | Amplify overhangs for knock-in of pRTS3/pRTS3.(+) |
| oPAS329 | TGAGCGAGGAAGCGGAAGAGCGCCCAATACcatcgcccagtcttctccc | Amplify overhangs for knock-in of pRTS3/pRTS3.(+) |
| oPAS270 | aggccaaataggccgttg | Amplify Tet(M) for pRTS3/pRTS3.(+) |
| oPAS227 | ctagtgaaatttccggctttattaaacttat | Amplify Tet(M) for pRTS3/pRTS3.(+) |
| oPAS267 | taaaaataagtttaataaagccggaaatttcactaggaagcaaacttaagagtgtg | Amplify ermB/ermB' for pRTS3/pRTS3.(+) |
| oPAS268 | atatacgcggcgcttaattttatggcctgaacattaaacacattccctttagtaa | Amplify ermB/ermB' for pRTS3/pRTS3.(+) |
| oPAS330 | cattaataagagagaggtactcattatgaattaca | Amplify knock-in DNA for integration into E. limosum from pRTS3, pRTS3.(+), pBgaBg, and LP |
| oPAS331 | catcgcccagtcttctcc | Amplify knock-in DNA for integration into E. limosum from pRTS3, pRTS3.(+), pBgaBg, and LP |
| oPAS620 | taagtgCAATTGAGAGGCGGTTTGCGTATTGG | Amplify pUC19 backbone for pRec-Cas1 |
| oPAS621 | tagctcTTAATTAAtaacactGCGGCCGCGACGAAAGGGCCTCGTGATA | Amplify pUC19 backbone for pRec-Cas1 |
| oPAS622 | aaggcagGCGCGCCattataGGTAACCggcataacagtgctgaata | Amplify RepH for pRec-Cas1 |
| oPAS623 | GCCTCTCAATTGcacttatcacattaagtatatactattattaaac | Amplify RepH for pRec-Cas1 |
| oPAS624 | agtgttaTTAATTAAgagctaccacAAAATGGAAAG | Amplify Tet(M) for pRec-Cas1 |
| oPAS625 | tatgccGGTTACCtataatGGCGCGCctgccttgattggaatagtttttag | Amplify Tet(M) for pRec-Cas1 |
| oPAS698 | TTGACTGCTAGCgcggtgtgaaataccgca | Amplify Cas9 for pRec-Cas2 |
| oPAS676 | tatgccTGTACAaaaaggccatccgtcaggatggccttctAATTAAGAGCTcggtacccc | Amplify Cas9 for pRec-Cas2 |
| oPAS696 | ccttttTGTACAggcataacagtgctgaatag | Amplify backbone from pRec-Cas1 for pRec-Cas2 |
| oPAS697 | caccgcGCTAGCAGTCAAAAGCCTCCGACCGGCTTTTGACTctgccttgattggaatagt | Amplify backbone from pRec-Cas1 for pRec-Cas2 |

| | | |
|---|---|---|
| oPAS682 | aaaacgaaaggctcagtcgaaagactgggcctttcgttttataccgcctttgagtgagct | Amplify pEL11 sgRNA for OE PCR for integration into either pCas-BgaB or pCas-p2TetO1 |
| oPAS683 | GTCAGAGGTTTTCACCGTCATCACCGAAACGCGcttatcatccccttttgcttatggagc | Amplify pEL11 sgRNA for OE PCR for integration into either pCas-BgaB or pCas-p2TetO1 |
| oPAS684 | tgtgcagctccataagcaaaaggggatgataagCGCGTTTCGGTGATGACG | Amplify target sgRNA for OE PCR for integration into either pCas-BgaB or pCas-p2TetO1 |
| oPAS685 | TTCTTTCCATTTTgtggtagctcTTAATTAACATTAATGCAGCTGGCACGAC | Amplify target sgRNA for OE PCR for integration into either pCas-BgaB or pCas-p2TetO1 |
| oPAS717 | cgcattcgtgaagcgattgaaaagtacaatgcaaagatatgctggatccttgacagctag | Amplify combined sgRNA casettes |
| oPAS718 | agtaaaattaatttcaatattccacaatattatattataagCTCACTCATTAGGCACCCC | Amplify combined sgRNA casettes |
| oPAS496 | ATGAAGCCGGCAAAGTTTAACGAGACCCTCCTCGCCtg | Amplify backbone from pRTS3.(+) for pBgaBg |
| oPAS497 | aaatgaaaccatattcgaaaaaatcgtctgtctgttatcctttcagaatttccagcagtt | Amplify backbone from pRTS3.(+) for pBgaBg |
| oPAS498 | tttcgaatatggtttcatttAAAATGGAAAGAAGCGCTGTTTTAAG | Amplify BgaB from pELIM2.2 for pGbaBg |
| oPAS499 | tgcaGGCGAGGAGGGTCTCGTTAAACTTTGCCGGCTTCATC | Amplify BgaB from pELIM2.2 for pGbaBg |
| oPAS340 | cgtttacgaaattggaacaggtaaagggcatttaacgacgaaactggctaaaataagtaaacaggtaacgtctattgaattagacagtcatctattcaacttatcgtcagaaaaattaaa | 120bp recombinase test oligo to repair ermB' in strain RTS (sense orientation) |
| oPAS341 | tttaatttttctgacgataagttgaatagatgactgtctaattcaatagacgttacctgtttacttattttagccagtttcgtcgttaaatgccctttacctgttccaatttcgtaaacg | 120bp recombinase test oligo to repair ermB' in strain RTS (antisense orientation) |
| oPAS486 | tgactgtctaattcaatagacgttacctgtttacttattttagccagtttcgtcgttaaa | 60bp antisense recombinagenic oligo for repairing ermB' into functional ermB |
| oPAS487 | atagacgttacctgtttacttattttagcc | 30bp antisense recombinagenic oligo for repairing ermB' into functional ermB |
| oPAS488 | tgactgtctaattcaatagacgttacctgcttgctaatcttggccagtttcgtcgttaaa | 60bp antisense recombinagenic oligo for repairing ermB' into functional ermB with wobble bases mutated to improve efficiency |
| oPAS489 | t*g*a*ctgtctaattcaatagacgttacctgcttgctaatcttggccagtttcgtcgtt*a*a*a | 60bp antisense recombinagenic s-oligo for repairing ermB' into functional ermB with wobble bases mutated to improve efficiency |
| oPAS490 | t*g*a*ctgtctaattcaatagacgttacctgtttacttattttagccagtttcgtcgtt*a*a*a | 60bp antisense recombinagenic s-oligo for |

| | | |
|---|---|---|
| | | repairing ermB' into functional ermB |
| oPAS518 | TATCGCGCCAGTTAACATGCGAGGTTACTTGCGCCATCAAAA TAAACGGTTGGCCTTTTC | 60bp antisense recombinegenic oligo for mutating BgaB to become inactive |
| oPAS549 | cctccttatttcaaattcaagtttatc | Colony PCR to confirm removal of pRec-Cas2 |
| oPAS679 | TGAATCTGTTTGTCAGCC | Colony PCR to confirm removal of pRec-Cas2 |
| oPAS349 | cctcctaacaacttaattatacccactattatta | Colony PCR to confirm removal of pEL11-BgaB |
| oPAS002 | CTACACCGAACTGAGATACC | Colony PCR to confirm removal of pEL11-BgaB |
| oEPS53 | ctaggtataatactagtattccctttaattttgtcaggttttagagctagaaatagcaag | Cloning pEL11-eps |
| oEPS54 | atttctagctctaaaacctgacaaaattaaagggaatactagtattatacctaggactga | Cloning pEL11-eps |
| oPAS763 | gcagtgctgttaatttttagcag | Colony PCR for clean eps knockout strain |
| oPAS764 | ggatgcctatatctttgaagatcaa | Colony PCR for clean eps knockout strain |
| oEPS55 | tataaaaaaagaacaccgcattgggtgttcttttttatggctataacttcgtataatgtatgct atacgaagttatatccttggaatctttttcattaacttctcccatgttctcaga | 120bp antisense recombinogenic oligo for eps operon knockout |

**Table S4:** sgRNA used in this study

| Target | Sequence | Description |
|---|---|---|
| pEL11 | aatttgagagggaacttagagttttagagctagaaat agcaagttaaaataaggctagtccg | Targets pEL11 for removal of recombinase plasmid. This sgRNA is encoded on pEL11-BgaB |
| BgaB | ACCGTTTATTTTGATGGAGCgttttagag ctagaaatagcaagttaaaataaggctagtccg | Targets unmutated BgaB to kill unrecombined cells. This sgRNA is encoded on pEL11-BgaB |
| eps operon | attccctttaattttgtcaggttttagagctagaaatagc aagttaaaataaggctagtccg | Targets unmutated eps operon to kill unrecombined cells. This sgRNA is encoded on pEL11-eps |

**Nucleotide sequence for *ermB'***

The sequence of three consecutive stop codons is highlighted. The original coding sequence in this region is: agtaaacag

ATGAACAAAAATATAAAATATTCTCAAAACTTTTTAACGAGTGAAAAAGTACTCAACCAAATAAT
AAAACAATTGAATTTAAAAGAAACCGATACCGTTTACGAAATTGGAACAGGTAAAGGGCATTT
AACGACGAAACTGGCTAAAATATGATAATAGGTAACGTCTATTGAATTAGACAGTCATCTATTC
AACTTATCGTCAGAAAAATTAAAACTGAATACTCGTGTCACTTTAATTCACCAAGATATTCTAC
AGTTTCAATTCCCTAACAAACAGAGGTATAAAATTGTTGGGAGTATTCCTTACCATTTAAGCAC
ACAAATTATTAAAAAAGTGGTTTTTGAAAGCCATGCGTCTGACATCTATCTGATTGTTGAAGA
AGGATTCTACAAGCGTACCTTGGATATTCACCGAACACTAGGGTTGCTCTTGCACACTCAAG
TCTCGATTCAGCAATTGCTTAAGCTGCCAGCGGAATGCTTTCATCCTAAACCAAAAGTAAAC
AGTGTCTTAATAAAACTTACCCGCCATACCACAGATGTTCCAGATAAATATTGGAAGCTATATA
CGTACTTTGTTTCAAAATGGGTCAATCGAGAATATCGTCAACTGTTTACTAAAAATCAGTTTCA
TCAAGCAATGAAACACGCCAAAGTAAACAATTTAAGTACCGTTACTTATGAGCAAGTATTGTC
TATTTTTAATAGTTATCTATTATTTAACGGGAGGAAATAA

**Nucleotide sequence for RecT from pEL11 (Refactored RecT from SA11 prophage)**

ATGGCAGTAAAGAACTCCCTGACAAAACAGGAGGCCAAGAAACCAACGTTCTCCAGCTATA
TCACGTCAGATGGCGTTAAGAGAAAAATTAACGAGATGGTAGGCGGAAAGGATGGCCAGAG
ATTTATCACGTCCATCATCTCCGCTGTGTCCACCAATAACGCGTTAGCGGCGTGCGACCAG
GGAACAATCTTAGCAGCAGCAATGTTGGGTGAGTCCTTAAAACTGTCTCCTTCTCCGCAGCT
TGGTCAGTATTATATGGTACCGTATGACCAGAAGGAGAAGAGAGATAAACAGGGCAATATTAT
CCAGAACGCTAAGAAAGTGGCTCAGTTCCAGCTTGGATACAAGGGCTACATCCAGCTCGCC
GAGAAATCCGGTCAGTATAAAAAACTTAATGTTGTTTCCATCAAGGAGGGTGAACTCGTAAA
GTTTGACCCGCTGAATGAAGAGATTGAAGTAAACCTGATCGAAGACGAAGAGGAGAGAGAA
GAAGCGAAGACGATCGGTTACTATGCAATGTTCGAGTATTTAAATGGATTTAGAAAGGCGATC
TATTGGTCCAAGAAAAAGATGGAGAAACACGCAGATCGTTACTCCAAGGCGTTTAATTTATC
CGTTTACGAAAAGATTAACGCAGGTGTCATCCCTGCCAAAGATCTGTGGAAGTATTCCTCTT
TTTGGTATAAGGACTTCGATGGCATGGCACACAAGACCATGCTGCGCCAGCTGATTTCAAAG
TGGGGAATCATGAGCATCGAGATGCAGACAGCGTTTGAGAAAGATATGACGGTAATCGACG
AAGATGGCAACGCAGAATACGTAGATAATATCGAATACGAGGCAGACAGCGAGTACATCGAG
ACAGCAGAAGTAGAAGAATTCAAGGAGGCAGCTATGACAGAGCACGAGGAGTCCGATGAT
GCGCAGATGACGATCCTGCCGAACGAGGCGGAGGATGAAGTCTTCAGCGAAGATGACTTC
TTCAACACGGAGGTATAA

**Nucleotide sequence from pEL11**

sgRNA binding region is highlighted in yellow and PAM site is highlighted in green.

TGCAGTCGAAGTGGGCAAGTTGAAAAATTCACAAAAATGTGGTATAATATCTTTGTTCATTAG
AGCGATAAACTTGAATTTGAGAGGGAACTTAGATGGTATTTGAAAAAATTGATAAAAATAGTT
GGAACAGAAAAGAGTATTTTGACCACTACTTTGCAAGTGTACCTTGTACTT

**Nucleotide sequence for BgaB**

sgRNA binding region is highlighted in yellow and PAM site is highlighted in green. Gray text shows where the recombinogenic oligo binds and the two red bases are the ones targeted for mutation by the oligo. In order, these are mutated A->C and G->A by the recombinase.

ATGAACGTTCTGTCCTCAATCTGCTACGGAGGAGATTACAACCCTGAGCAATGGCCAGAGG
AAATTTGGTATGAAGACGCTAAGTTGATGCAAAAAGCGGGGGTGAATTTAGTGTCTTTAGGG
ATTTTCAGTTGGAGCAAAATCGAACCGTCTGATGGAGTGTTCGACTTTGAATGGCTCGACAA
GGTTATAGACATACTATATGACCACGGTGTGTATATCAACTTGGGGACGGCGACCGCAACCA
CTCCGGCTTGGTTTGTAAAAAAATATCCGGATTCTTTGCCGATCGATGAAAGCGGAGTCATT
CTCTCGTTTGGCAGCCGCCAACATTATTGTCCTAATCATCCTCAATTAATTACGCACATAAAG
AGACTTGTGAGGGCTATTGCCGAACGGTATAAAAATCATCCGGCACTCAAAATGTGGCATGT
TAACAATGAGTATGCATGTCACGTTTCCAAATGTTTTTGCGAGAATTGCGCCGTCGCGTTTC
GGAAGTGGCTAAAGGAAAGATATAAAACAATCGACGAACTTAATGAACGTTGGGGTACAAAC
TTTTGGGGACAGCGATACAACCATTGGGATGAAATCAATCCCCCTAGAAAGGCACCGACGT
TTATCAATCCAAGCCAGGAACTTGACTACTACCGTTTTATGAATGACTCAATTCTCAAGTTGT
TTTTAACAGAAAAAGAAATTTTACGCGAGGTAACACCAGATATTCCGGTGTCAACTAATTTCA
TGGGTTCATTCAAACCGCTGAACTATTTTCAATGGGCGCAGCATGTAGATATTGTGACATGG
GACTCATATCCTGACCCCAGAGAGGGCCTTCCGATCCAGCACGCCATGATGAATGACCTTAT
GCGTTCTCTTAGAAAAGGCCAACCGTTTATTTTGATGGAGCAGGTAACCTCGCATGTTAACT
GGCGCGATATTAATGTTCCAAAACCGCCAGGTGTAATGCGTTTATGGAGTTATGCAACGATT
GCCCGTGGCGCCGATGGTATTATGTTTTTCCAGTGGCGGCAAAGTAGAGCAGGAGCTGAAA
AATTCCACGGTGCAATGGTGCCTCACTTTTTGAACGAGAATAATAGAATTTATCGCGAAGTTA
CACAGCTTGGACAAGAGCTGAAAAAGCTGGATTGTTTGGTCGGATCTAGAATCAAGGCCGA
GGTCGCGATCATCTTTGATTGGGAAAACTGGTGGGCTGTCGAACTGTCCTCCAAACCGCAT
AACAAACTGCGCTATATTCCTATAGTTGAAGCTTATTACAGGGAATTATATAAACGGAATATTG
CTGTCGATTTTGTCCGCCCATCTGATGATCTAACAAAATACAAAGTGGTTATCGCCCCAATGT
TATATATGGTCAAAGAGGGAGAAGATGAAAACCTTCGGCAATTTGTTGCGAACGGCGGCACT
CTGATTGTCAGCTTCTTCTCGGTCATCGTCGACGAAAATGACCGAGTACATCTCGGCGGATA
TCCTGGCCCTCTGCGAGATATTTTGGGCATCTTTGTTGAGGAATTTGTACCTTACCCGGAAA
CCAAAGTAAACAAAATCTATAGCAACGATGGCGAATATGATTGCACGACGTGGGCGGACATA
ATCCGGTTAGAAGGGGCAGAACCTCTCGCGACATTTAAGGGGGATTGGTATGCAGGACTTC
CGGCGGTTACACGTAACTGCTACGGTAAAGGAGAGGGGATCTACGTCGGTACGTATCCGGA
TAGCAATTATTTAGGCAGGCTTTTAGAACAGGTCTTCGCTAAACATCATATTAACCCCATTCTT
GAAGTAGCTGAAAATGTAGAGGTGCAGCAAAGAGAGACTGATGAATGGAAGTATCTTATTAT
TATCAATCATAATGATTACGAAGTGACACTGTCACTGCCGGAAGATAAAATATACCAGAATATG
ATTGATGGGAAATGTTTTCGAGGAGGCGAACTGAGGATTCAAGGCGTTGATGTGGCAGTGC
TGCGCGAGCATGATGAAGCCGGCAAAGTTTAA
